# Developmental system drift in the patterning of the arthropod tarsus

**DOI:** 10.1101/2025.07.08.663771

**Authors:** Benjamin C. Klementz, Sophie M. Neu, Ethan M. Laumer, Emily V.W. Setton, Isaac A. Hinne, Austen A. Barnett, Max Hämmerle, Georg Brenneis, Monika Gulia-Nuss, Prashant P. Sharma

## Abstract

The current understanding of proximodistal axis patterning in arthropod legs is grounded in insect models. The paradigm for appendage evolution in this phylum is that the gene regulatory network responsible for leg subdivision and patterning is broadly conserved. Recent surveys of these genes have suggested that chelicerate exemplars exhibit divergent appendage patterning dynamics, though functional data remain limited. One salient mismatch in expression occurs in homologs of the homeobox gene clawless. In insects, clawless is expressed in the distalmost leg territory, specifying the claw-bearing pretarsus. In the harvestman, Phalangium opilio, clawless occupies a broad tarsal domain early in development, localizing later to the metatarsus-tarsus boundary, suggestive of a tarsal patterning function. Here, we tested the function of harvestman clawless using RNAi. Unlike insects, we show that clawless knockdown results in disrupted tarsal growth and patterning of its proximal segmental boundary, with no effect on the claw. Truncation of the tarsus is associated with defective tarsomere formation. We additionally surveyed clawless homologs in exemplars of chelicerate diversity, which suggests that the tarsal-patterning function for clawless was likely present in the chelicerate common ancestor. These results, alongside available expression data, suggest panarthropod appendage patterning exhibits numerous cases of developmental system drift.

## Introduction

The hyperdiversity of extant arthropods is partly attributed to the modularity and adaptability of their segmented appendages. Almost every segment of the arthropod walking leg has been modified and outfitted with evolutionary novelties across the span of arthropod diversity. Further modifications of the walking leg developmental program are understood to underlie the diversification of appendage types, such as insect mandibles, crustacean pleopods, and sexually dimorphic gonopods in millipedes [1,2].

Much of what is known about appendage patterning in arthropods stems from seminal works in insect models such as the fruit fly Drosophila melanogaster and the red flour beetle Tribolium castaneum, which share the five-segmented condition archetypal of insects. Work on satellite model systems has elucidated evolutionary dynamics of leg patterning, such as the role of different signaling pathways in leg segment morphogenesis. Such datasets have broadly supported the interpretation that leg patterning is conserved across the phylum, with respect to (1) the identity and spatial arrangement of transcription factor domains that establish the proximo-distal axis, (2) the involvement of Notch-Delta signaling in establishing segment boundaries, and, to a lesser extent, (3) the identity of downstream genes that further subdivide the developing leg.

Drawing upon this pattern of conservation, some workers have proposed to align leg segments across the phylum through the lens of gene expression and functional data, with the goal of reconstructing the ancestral arthropod leg [3]. Our recent work, however, has highlighted the variability of leg patterning and gene expression dynamics across Chelicerata, the subdivision of arthropods that includes sea spiders and arachnids.

Chelicerates are particularly notable for the anatomical variability of their appendage segment complement across extant orders, resulting in historical discord over their leg segment nomenclature. We were able to show that an intermediate leg segment, the patella, is established via a new expression domain of the homeobox gene extradenticle in a taxon-specific manner, with disruption of exd expression resulting in loss of the distal patellar boundary in a daddy-longlegs [4]. As an ancillary investigation, we also showed that many transcription factors known to pattern specific territories of the fruit fly leg exhibited markedly different expression domains in the developing legs of a daddy-longlegs and a sea spider [5].

The most curious of these divergent expression domains pertain to the transcription factors clawless (cll) and aristaless (al) (figure 1a). In the fruit fly, these two genes are required for the patterning of the tarsal claws of the legs and of the arista of the antenna, and are co-expressed in the Distal-less-positive region of the imaginal disc that corresponds to the distal tip. The claw is initially specified by activation of the homeobox transcription factors BarH1 and BarH2 (collectively, Bar), aristaless, clawless, and Lim1 as a product of distal-to-proximal epidermal growth factor receptor (EGFR) signaling gradients [6–8]. Al and Cll proteins bind cooperatively, likely to sites identified in the putative Bar enhancer, to repress Bar in the pretarsus, initially specifying the boundary between distal tarsus and proximal pretarsus. Likewise, Bar represses Lim1 in the distal tarsus, itself necessary for further activation of aristaless and clawless, thus forming a regulatory feedback loop that leads to complete separation of tarsus and pretarsus [6,7,9–11]. Loss-of-function mutants for either gene exhibit a comparable phenotype, with losses of pretarsi and antennal aristae.

**Figure 1.**
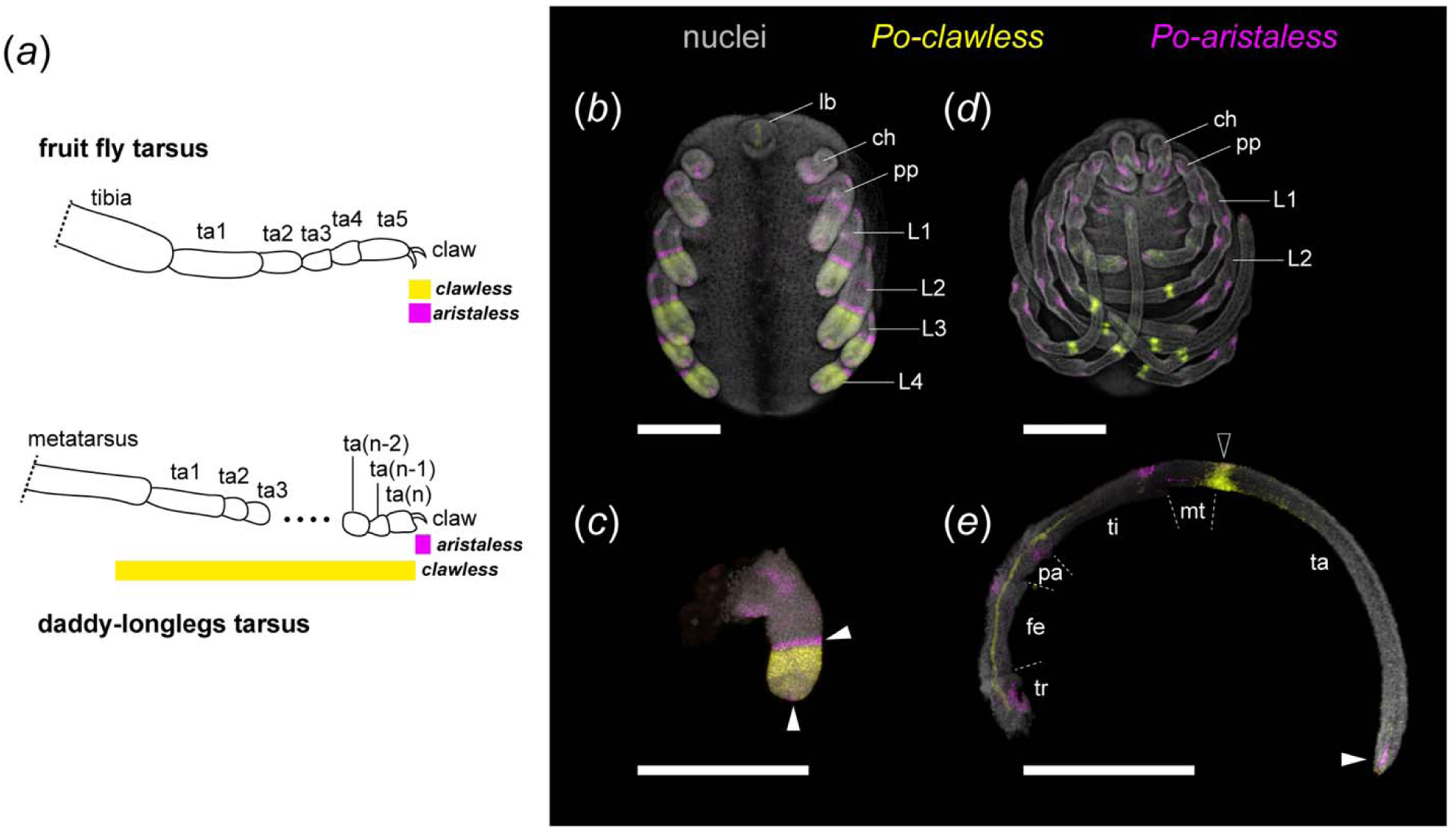
Divergent dynamics of distal limb patterning gene expression between arthropod taxa. (a) Schematic representations of cll and al expression domains in the distalmost podomeres of the fruit fly and daddy-longlegs walking leg. (b-e) Multiplexed expression of Po-cll (yellow) and Po-al (magenta) with nuclear counterstaining (grey) in P. opilio embryos. (b) Stage 10 embryo. (c) Leg II of stage 10 embryo. Note the proximal expansion of Po-cll beyond al-positive cells in the distal terminus. (d) Stage 13 embryo. (e) Leg II of stage 13 embryo. Note the late localization of Po-cll to the metatarsal-tarsal boundary (black arrowhead) and depletion of tarsal expression. White arrowheads: al-positive expression domains. The proximal domain adjacent to Po-cll disappears by stage 13. Scale bars: 100 µm. Abbreviations: ch – chelicerae; fe – femur; L – leg; lb – labrum; mt – metatarsus; pa – patella; ta – tarsus; ti – tibia; tr – trochanter.

In marked contrast, in the daddy-longlegs, we found that aristaless retains a limited expression domain in the tips of the walking legs (comparable to insects), whereas clawless was broadly expressed throughout the distal limb bud in early stages of leg elongation, but absent from the distal-most tip of the territory marked by aristaless-positive cells (figure 1b-c) [5]. Later in development, when podomere boundaries are visible, clawless spans the distal metatarsus and proximal tarsus, with the highest intensity of expression at the segmental boundary (figure 1d-e). Similar expression domains were likewise observed in the appendages of the sea spider Pycnogonum litorale, supporting the inference that the dynamics of these genes are not restricted to daddy-longlegs [5].

The evolutionary divergence of distal patterning in insect and chelicerate legs is unexpected because the homology of the distal-most territory of the walking leg has never been questioned, nor have functional data (where available) ever suggested deviation of tarsal patterning across Arthropoda. The distal region of the arthropod appendage, the tarsal claw, seems to be retained across all arthropod taxa, despite variations in nomenclature. In terrestrial lineages, single or paired tarsal claws (e.g., unguis) are borne on the distal terminus, the pretarsus (“dactylus” of crustaceans; “main claw” or “dactylus” of sea spiders), differentiated from true podomeres by the lack of intrinsic musculature (note: the pretarsal claws are commonly referred to as the tarsal claws in taxonomic literature of many arachnid groups; for the purpose of simplicity, we use the term tarsal claw here). However, in arachnids the pretarsus is reduced to a ring-like sclerite operated by a single pair of antagonistic depressor and levator muscles [12]. The pretarsus can bear morphological modifications to improve traction reinforcement, such as the pulvillus or arolium, which facilitate adherence to smooth substrates. In spiders, additional claws serve the function of silk-handling hooks.

Here, we assessed the function of clawless in the daddy-longlegs Phalangium opilio using a gene silencing approach. Consistent with its expression pattern, we show that depletion of clawless yields a fusion of metatarsus and tarsus, with truncation of the latter and loss of tarsomeres (articles of the tarsal segment), but has no effect on the tarsal claw. To assess the evolutionary dynamics of this gene, we surveyed clawless homologs in an array of chelicerate models. Polarizing these dynamics using data from insect, myriapod, onychophoran, and tardigrade counterparts, we show that the acquisition of a tarsal selector function for clawless evolved in the last common ancestor of Chelicerata, highlighting a case of developmental systems drift in arthropod appendage patterning.

## Material and Methods

### (a) Bioinformatics and orthology inference

Homologs of clawless and aristaless were identified in the genomes of the chelicerates Phalangium opilio (Opiliones) [13], Parasteatoda tepidariorum (Araneae) [14], Ixodes scapularis (Parasitiformes), [15] Archegozetes longisetosus (Acariformes) [16], and Pycnogonum litorale (Pycnogonida) [17]. As queries, we used the peptide sequences of these two genes from the velvet worm Euperipatoides kanangrensis (GenBank accession nos. CDK60407.1; CDK60408.1) and performed tBLASTn searches against the annotated CDS files of each genome assembly, retaining hits with e-value < 1×10^-20^. Sequence identities were validated using SMART-BLAST, followed by multiple sequence alignment with CLUSTAL Omega [18]. Gene trees were inferred using IQ-TREE v. 1.6.12 [19] with an LG+I+G substitution model and nodal support was estimated using 1000 ultrafast bootstrap replicates. Alignments and gene trees are provided in electronic supplementary material files S1-S4.

### (b) Gene expression assays

Embryos were fixed and assayed for fluorescent detection of gene expression following established or minimally modified protocols, as detailed previously for chelicerate species [4]. For I. scapularis culture, all procedures involving animal subjects were approved by the Institutional Animal Care and Use Committee (IACUC) at the University of Nevada-Reno (IACUC #21-01-1118-1). In species other than A. longisetosus, we targeted selected developmental stages where podomeres could be individually recognized.

Probe design for hybridization chain reaction (HCR) consisted of 6-20 probe pairs, depending upon the length of the available template sequence. Probes were designed either with the HCR Probe Maker tool [20], or via submission of CDS to Molecular Instruments.

Input CDS templates for proprietary Molecular Instrument probe design and probe sequences obtained from the HCR Probe Maker tool for all species are provided in electronic supplementary material file S5.

Due to the challenges of autofluorescence of spider yolk, we deployed single-channel colorimetric expression assays for P. tepidariorum [21]. Gene cloning, probe synthesis, and traditional colorimetric in situ hybridization followed the approaches detailed in our previous studies of this species [22,23]. Primers for design of RNA antisense probes are also provided in electronic supplementary material file S5.

### (c) RNA interference

Fragments of Po-cll were amplified using standard PCR protocols and cloned using a TOPO TA Cloning Kit using One Shot Top10 chemically competent Escherichia coli (ThermoFisher) following the manufacturer’s protocol, and PCR product identity was verified via Sanger sequencing with M13 universal primers. Double-stranded RNA (dsRNA) was synthesized following the manufacturer’s protocol using a MEGAscript T7 kit (Ambion/Life Technologies) from amplified PCR product. The quality of dsRNA was assessed and concentrations adjusted using a NanoDrop ONE to 3.8-4.2 µg/µL. dsRNA was mixed with vital dyes for visualization of injections. Microinjection under halocarbon-700 oil (Sigma-Aldrich) was performed as previously described [24]. Subsets of developing embryos were fixed for HCR; the remainder was developed at 26°C until hatching.

### (d) Imaging

Brightfield microscopy was performed using a Nikon SMZ fluorescence stereomicroscope mounted with a DSFi2 digital color camera driven by Nikon Elements software. Images of P. opilio appendages were captured at varying focal planes and compiled into focused stacks with Helicon Focus v. 6.6.1. Confocal laser scanning microscopy was performed using a Zeiss LSM 710 or a Zeiss LSM 980 microscope driven by Zen software.

For P. litorale, confocal laser scanning microscopy was performed with a Leica SP5 microscope, driven by LAS-AF software. Beyond the documentation of gene expression (594nm and 633nm laser lines) and Dapi counterstain (405nm laser line), cuticular autofluorescence was separately recorded with the 488nm laser line. Using the software Amira 3D (version 2021.1; ThermoFisher Scientific), the cuticular signal in the 488nm channel was semi-automatically segmented (grey-value based thresholding) and the voxels included in the resulting material were set to grey value 0 in all other channels via the “Arithmetic” function, resulting in the separation of cuticular autofluorescence from gene expression signals.

## Results and Discussion

### (a) clawless is required for the patterning of the harvestman tarsus, but not the claw

Consistent with the late localization of Po-cll expression to the metatarsus-tarsus boundary, Po-cll RNAi yielded phenotypes exhibiting fusion of the metatarsus and tarsus (figure 2).

**Figure 2.**
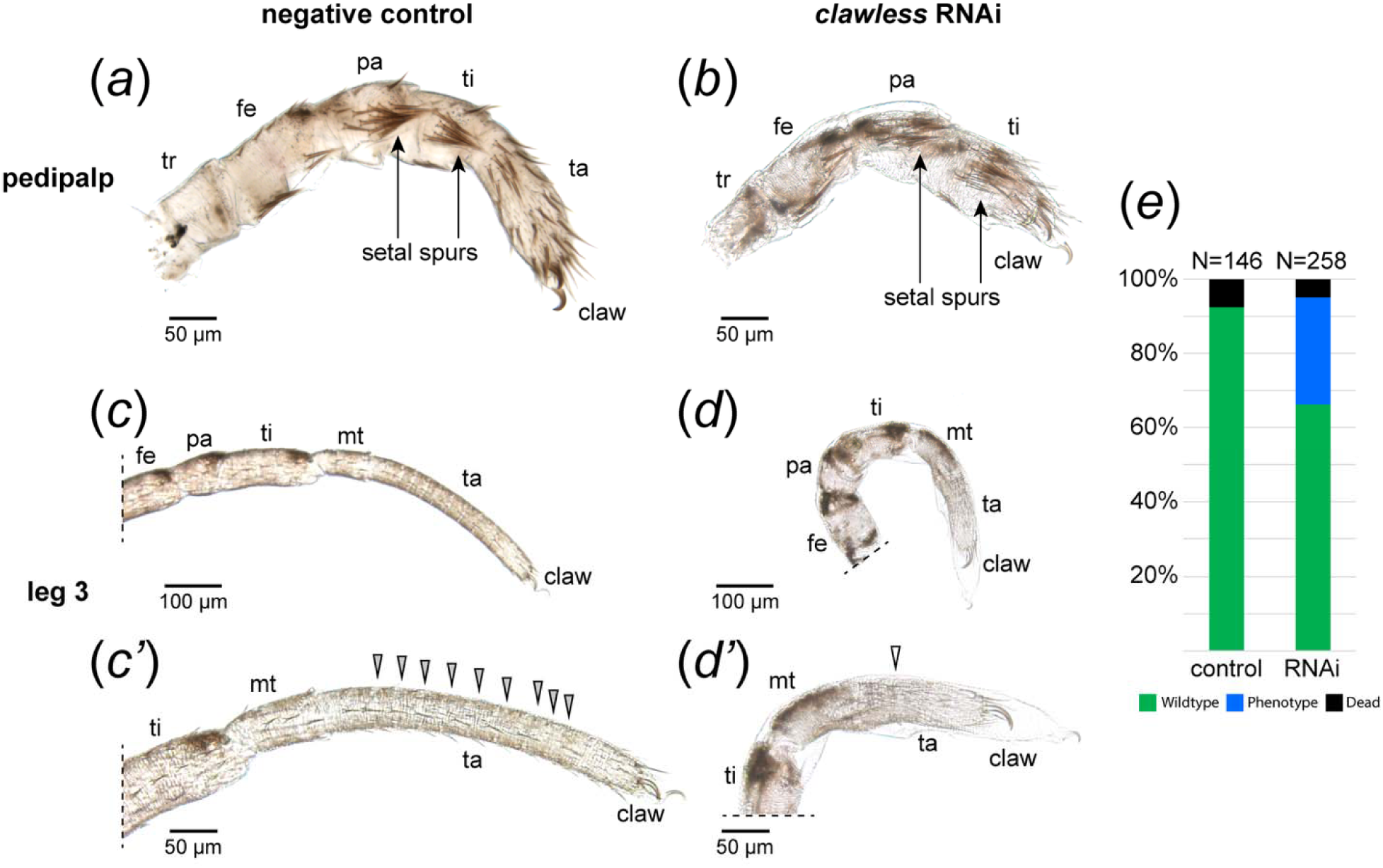
Po-cll is necessary for the development of the tarsus, but not the claw, in P. opilio. (a-d) Appendage mounts of hatchlings in lateral view. (a) Pedipalp of negative control hatchling. (b) Pedipalp of Po-cll RNAi hatchling. (c) Leg III of negative control hatchling. (c’) Magnification of tarsus III, showing tarsomeres (grey arrowheads). (d) Leg III of Po-cll RNAi hatchling. (d’) Magnification of tarsus III in Po-cll RNAi hatchling; note truncation of the tarsus and absence of tarsomeres (white arrowhead). (e) Distribution of phenotypic outcomes in negative control and Po-cll RNAi experiments.

Following embryogenesis, 28.7% (n = 74/258) of hatchlings exhibited the fusion phenotype, with slight variations in the resulting deformation (figure 2e). Mild phenotypes were characterized by an outward bulging of the tissue at the presumptive malformed segmental boundary during embryonic stages (figure 2c-d’; 3a-b), whereas severe phenotypes were characterized instead by distal appendages bearing a pronounced bent or kinked appearance, which we attribute to asymmetric projections of tissue at the malformed joint (figure S1). The severe phenotype, however, never impacted all appendages in a single embryo; one or two appendages bearing the kinked appearance were common. Both phenotypic classes, however, are characteristic of podomeric fusions following RNAi against limb-patterning transcription factors in both arachnid and insect exemplars [4,25–27]. No Po-cll RNAi hatchlings exhibited defects in or absence of the tarsal claws or of cheliceral tips (contra insects, [6]).

Coincident with the segmental fusion was a significant shortening of the tarsus in Po-cll RNAi hatchlings, in tandem with a reduction in the number of correctly formed tarsomeres (figure 2c’,d’). This truncation is most dramatically observed in the pedipalp, an appendage that is distinguished from the walking legs by the lack of a metatarsus. In the palps of wildtype hatchlings, two prominent setal tufts (“spurs”) occur in the distal territories of the patella and tibia (figure 2a). In Po-cll RNAi hatchlings, however, the tibial spur appears proximally adjacent to the claws, suggesting near complete absence of the tarsus, without affecting the tibia (figure 2b). This result supports the interpretation that Po-cll is required specifically for maintenance of the tarsal segment in both the leg and the pedipalp.

The effect of Po-cll RNAi on the tarsomeres is reminiscent of failed tarsomere formation in knockdowns of EgfrA [13]. In insects, simultaneous formation of tarsomeres is achieved by establishment of a distal-to-proximal gradient of EGFR activity, which provides positional information required for the precise activation of transcription factors like dachshund, bric-a-brac, and Bar; these genes are responsible for subdivision of the tarsus into proximal, medial, and distal domains, respectively. Regulatory interactions amongst these and other transcription factors result in unique gene expression combinations in each of the five tarsomeres of the fruit fly. Thereafter, heterogeneous expression of Notch and its ligands Delta and Serrate promote regions of cell proliferation via activation of downstream targets like AP-2, and regions of apoptotic activity via activation of targets reaper and head involution defective, resulting in the formation of tarsomere boundaries [8,28,29]. By contrast, in P. opilio tarsomeres are not formed simultaneously, but rather sequentially as the tarsus elongates in later stages of development. To characterize the nature of tarsomere patterning in P. opilio, we first characterized the expression dynamics of Po-Delta (Po-Dl) and Po-Serrate (Po-Ser) in the elongating tarsus.

In stage 9 embryos, Po-Ser is expressed in ring domains corresponding to putative segmental boundaries, including the metatarsal-tarsal boundary (figure S2a). Expression in this stage is strongest in the proximal boundaries, and only weakly detected in distal appendage territories. In later stages, Po-Ser remains localized to segmental boundaries, but does not exhibit additional ring domains in the elongating tarsus, likely refuting a direct role in tarsomere formation (figure S2b-d). Tarsal expression of Po-Ser is diffuse and patchy, likely coincident with cells fated for sensory or neural fates given previously characterized roles for Notch signaling in the regulation of proneural genes including the achaete-scute and Enhancer of split complexes [30,31].

Po-Dl likewise localizes to putative segmental boundaries during early stages of appendage elongation, again including the metatarsus-tarsus boundary. Yet by stage 9, an additional stronger domain of Po-Dl expression is detected in the leg II tarsus, distal to the segmental boundary of Po-Ser expression (figure S2e). In progressively later stages, the same ring domain appears in legs I, III, and IV. In these stages, the initially singular domain in each appendage multiplies, with sequential addition of ring domains distally (figure S2b-g).

Given the sequential appearance of these domains, as well as the canonical role of Delta in patterning segmental boundaries, Po-Dl appears to be a reliable readout of sequential tarsomere formation in P. opilio. Consistent with this interpretation, such ring domains are absent in the pedipalpal tarsus, which does not develop tarsomeres. Earlier onset of tarsal expression in leg II likewise coincides with observations of higher tarsomere counts in leg II, owing to its antenniform morphology (elongation of podomeres; concentration of sensory organs), analogous to the antenniform appendages of lineages such as Amblypygi and Thelyphonida.

In Po-cll RNAi embryos, no ring domain of Po-Dl is detected in the metatarsal-tarsal boundary, and no additional ring domains are observed in the tarsus (figure 3a’-d). Distal expression is restricted to putative sensory structures. The disruption of Po-Dl expression dynamics in this region is strongly suggestive of failed tarsomere formation, which is also observed in the tarsi of hatchlings.

**Figure 3.**
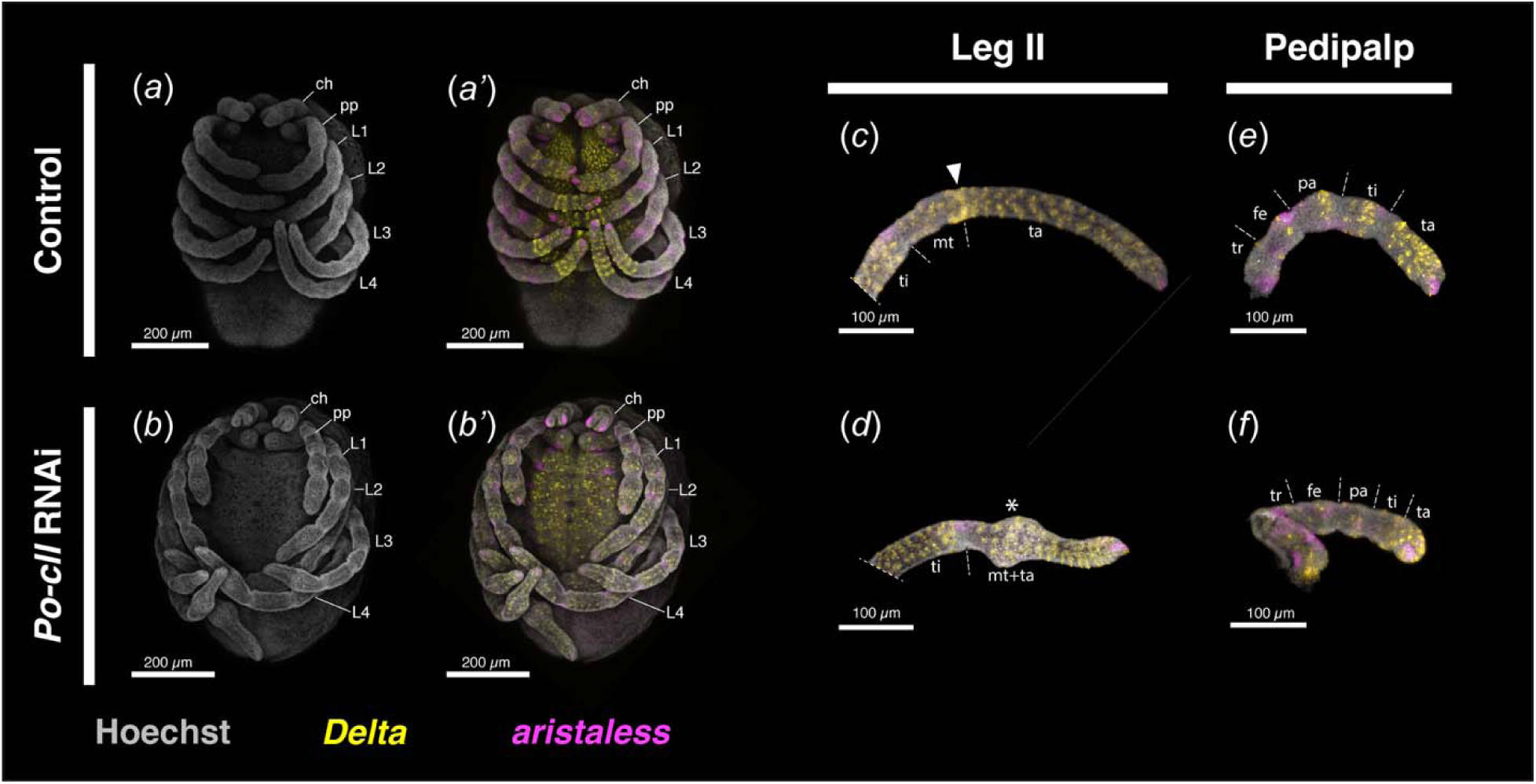
Knockdown of Po-cll disrupts tarsomere patterning mechanisms. (a, b) Stage 11 embryos with isolated nuclear counterstaining (grey). (a’-f) Multiplexed visualization of nuclear counterstaining, and expression of Po-Dl (yellow) and Po-al (magenta) in control and Po-cll RNAi treatments. (a’) Same embryo as in (a). (b’) Same embryo as in (b). (c) Leg II of negative control embryo. Note the strong ring of Po-cll at the metatarsus-tarsus boundary (arrowhead). (d) Leg II of Po-cll RNAi embryo. Note absence of the Po-cll ring (asterisk) and truncation of the tarsus. (e) Pedipalp of control embryo. (f) Pedipalp of Po-cll RNAi embryo. Abbreviations as in Figure 1.

Taken together, these results support a novel role for Po-cll in the patterning of the tarsus, rather than the distalmost territory (claws; aristae) as in other arthropod lineages.

### (b) A broad tarsal domain of clawless is phylotypic of Chelicerata

To pinpoint the phylogenetic origin of these expression dynamics, we surveyed the expression of cll in phylogenetically significant representatives of Chelicerata, including the sister group to the remaining chelicerates (the sea spiders). Where available, al orthologs were also surveyed to test the association of al expression with the claw.

In the acariform mite, Archegozetes longisetosus (Acariformes: Sarcoptiformes), Al-cll expression appears early in the appendage primordia (figure S3a-a’’’), localizing in limb bud stages to a cap-like distal domain (figure 4a, S3b-e’’’). Unlike the other surveyed taxa, however, Al-cll does not exhibit heterogeneous intensity; no clear ring domains were observed. The onset of Al-al expression occurs only after initial outgrowth of the limb buds from the body wall (figure S3b-b’’’). Al-al is expressed in two distinct domains, one as a diffuse domain in the proximal territory of all five appendage pairs of the hexapodous embryo, as well as a minute domain of several al-positive cells in the distal terminus within the broader domain of Al-cll (figure 4a, S3c-e’’’). Despite the lack of clear podomere boundaries in the stages examined, the extension of Al-cll expression proximally beyond the distal Al-al domain reflects the same expression dynamics observed in P. opilio.

**Figure 4.**
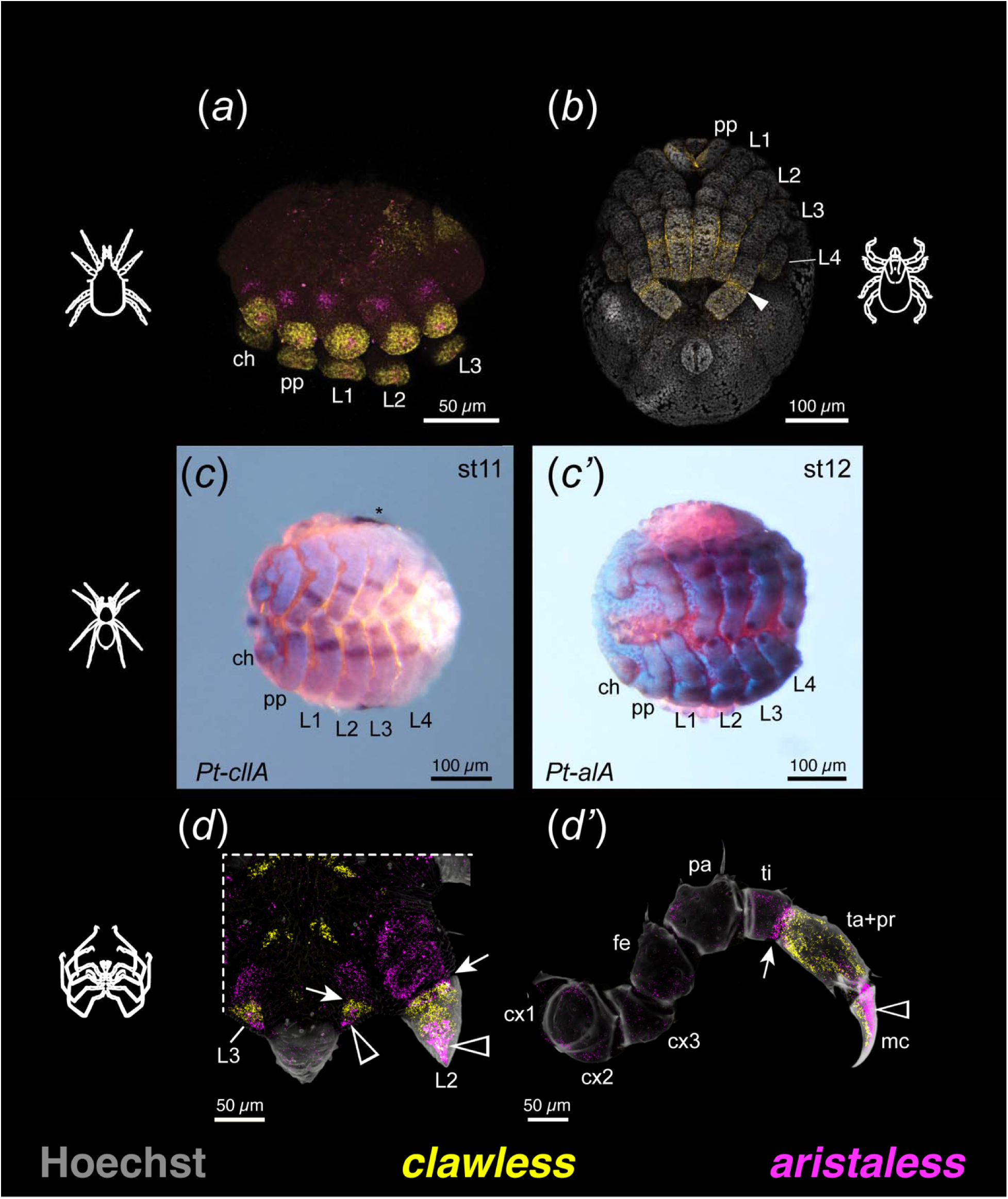
Proximal expansion of cll expression beyond the distal terminus is phylotypic of Chelicerata. (a) Multiplexed expression of cll (yellow) and al (magenta) in the acariform mite A. longisetosus. (b) Stage 13 embryo of the parasitiform tick I. scapularis with expression of cll. White arrowhead: basi- and telotarsus segmental boundary. (c) Stage 12 embryo of the spider P. tepidariorum stained for expression of cllA with nuclear counterstaining. (c’) Stage 11 embryo of P. tepidariorum stained for expression of al with nuclear counterstaining. Asterisk denotes non-specific stain in the yolk. (d-d’) Expression of Pl-cll and Pl-al with cuticular autofluorescence (grey) in the sea spider P. litorale. (d) Ventral view of instar IV posterior body pole and leg 2 limb bud. (d’) Leg 1 of advanced instar V. Black arrowhead: ectal Pl-al domain in the distal limb bud tips and in the main claw of legs. White arrowhead: mesal Pl-cll domain in the distal limb bud tips and in the main claw of legs. Arrow: proximal boundary of Pl-cll expression in limb buds and region of Pl-cll and Pl-al co-expression at the tibial-tarsal boundary in developing legs. Abbreviations: cx – coxa; mc – main claw; pr – propodus; others as in Figure 1.

Embryos of the parasitiform tick, Ixodes scapularis (Parasitiformes: Ixodida), were surveyed for Is-cll expression at stage 13 wherein podomere boundaries are visible in the first three pairs of walking legs; only the distal tip of the fourth pair of legs are visible during retraction and formation of the hexapodous larva [32]. I. scapularis embryos were also surveyed for Is-al, but expression was not detected in the stages surveyed (data not shown). At this stage, Is-cll is expressed broadly throughout the distal appendage territories of the walking legs, including weakly in the remaining leg IV bud (figure 4b). As in the harvestman, however, expression in the legs is heterogeneous; the highest intensity of expression localizes to the putative boundary of basi- and telotarsus. This pattern is suggestive of shared distal appendage patterning dynamics with the harvestman. Likewise, this expression domain is consistent with homology of the tick basitarsus and harvestman metatarsus, supporting the interchangeable use of both metatarsus and basitarsus by some authors [33,34].

The spider Parasteatoda tepidariorum, as a member of Arachnopulmonata (a group of six chelicerate orders ancestrally bearing book lungs), bears a whole genome duplication [35]. Two copies of clawless and one of aristaless were retrieved from the genome assembly; we were able to obtain expression data for one cll copy and the al copy, but the available data substantiate a chelicerate-specific pattern. The expression of Pt-cllA begins with a broad distal, cap-like domain in the early limb buds as they begin outgrowth (figure S4a-e). By stage 12, Pt-cllA expression is notably heterogeneous, with only weak signal in the distalmost territories, but as in both the harvestman and tick there is a stronger ring domain localized to the putative metatarsus-tarsus boundary (figure 4c, S4f). Pt-al expression likewise initiates shortly after emergence of the limb buds, appearing as two distinct domains, one a broad proximal domain near the appendage base, and a small region of al-positive cells in the distal terminus (figure S5a). In slightly later stages, an additional third region of strong expression is detected in the medial limb bud (figure S5b-c). In stages when podomere boundaries are visible, Pt-al is detected in four patch domains, one at the appendage base, one in the femur, one in the tibia, and one in the distal terminus expression (figure 4c’, S5e). Crucially, these domains are restricted only to the ectal portion of the appendages. In contrast to Pt-cllA, Pt-al is additionally detected in patchy, ectal domains of the tubular tracheae, book lungs, anterior and posterior spinnerets (figure S5d,f).

The sea spider Pycnogonum litorale hatches as a protonymphon larva with only three appendage pairs: the chelicerae and two pairs of larval appendages (mostly resorbed in adulthood in Pycnogonidae). The walking legs I-IV are added sequentially alongside the remaining body segments during successive molts. We previously showed that Pl-cll is strongly associated with two distal podomeres (the tarsus and propodus), but the exact homology of the sea spider claw (sometimes termed dactylus) is an outstanding question and impedes unambiguous comparison [5]. To surmount this ambiguity, we examined embryos for co-expression of Pl-cll and Pl-al. In instar II, both Pl-cll and Pl-al are detected in the leg I primordium that flanks the posterior body pole. Both genes are partially co-expressed at the distal-most area of the primordium; the more restricted Pl-al territory is embedded in the wider cap-like Pl-cll expression domain (figure S6a). By instar III, when the leg I bud has begun outgrowth from the body wall, Pl-al is weakly expressed in a wide domain in the proximal limb bud, as well as strongly at the distal terminus (figure S6b,c; S7a). Pl-cll expression occupies the intervening medial appendage domain, with a sharp proximal border at the boundary of Pl-al expression. At the leg’s distal terminus, the Pl-cll domain extends into the mesal compartment of the developing claw, complementary to the ectally concentrated Pl-al territory (figure 4d, S7a,a’,b’). In instars IV and V, wherein the first functional podomeres have formed in legs I and II, respectively, Pl-cll is expressed in the precursor podomere that will eventually divide into mature tarsus and propodus, and the mesal compartment of the claw (figure 4d’, S7c-d’). Pl-al is detected diffusely in all further proximal podomeres, as well as strongly in the ectal compartment of the claw (figure S7c-d’). Both genes exhibit little to no co-expression, with the exception of a narrow ring of elevated Pl-al expression at the proximal boundary of the Pl-cll domain (figure S7d,d’). The leg’s short tarsus will differentiate in this co-expression domain which becomes more distinct as the leg’s growing tissues condense prior to the molts. Once the tarsus and propodus are fully separated (instar VI), the ring of Pl-cll and Pl-al co-expression becomes more diffuse; only a small area with distinct expression overlap persists in the ectal joint of the tarsus (figure S7e,e’). By contrast, the claw retains its ectal Pl-al and mesal Pl-dll domains (figure S7e,e’). These results support the interpretation that the sea spider tarsus, propodus, and claw are homologs of the arachnid metatarsus, tarsus, and pretarsus, respectively. This reconstruction suggests that the additional (auxiliary) claws observed in some sea spider species may represent further derivations of the pretarsus, comparable to accessory claws (e.g., spider silk hooks) or pseudonychia of arthropods.

Taken together, these data support the interpretation that a broad cll domain subtending the pretarsus (marked by al) is phylotypic for Chelicerata (figure 5). This pattern stands in opposition to the conserved cll dynamics seen in the mandibulate arthropods. In the holometabolous insect models Drosophila melanogaster and Tribolium castaneum, cll expression is restricted to the center of the imaginal disc or the pretarsus of the leg, respectively [6,36,37]. Likewise, in the pill millipede Glomeris marginata, cll is again restricted to the distal terminus of the developing walking legs [38]. These dynamics are also characteristic of Panarthropoda broadly. In the lobopods of onychophorans, the sister group to the arthropods, cll is expressed in the distal tip, coincident with the onset of al expression in the same territory [39]. The lobopods of tardigrades also bear distal expression of cll [40]. In this lineage, cll expression is also detected proximal to the distal tip, extending to the medial appendage territory. However, unlike the proximal expansion of cll in chelicerates, proximal tardigrade cll is only detected in the posterior compartment of the appendage. The absence of cll expression at stage 17, wherein the claws are fully formed, is suggestive of a primary role in claw formation. Unlike other panarthropods, tardigrade al expression dynamics seem to be unique. While expressed in the distal appendage of the tardigrade, al is restricted only to an internal compartment, absent from the distal ectoderm, with only limited co-expression with cll at the margin of the ectoderm.

**Figure 5.**
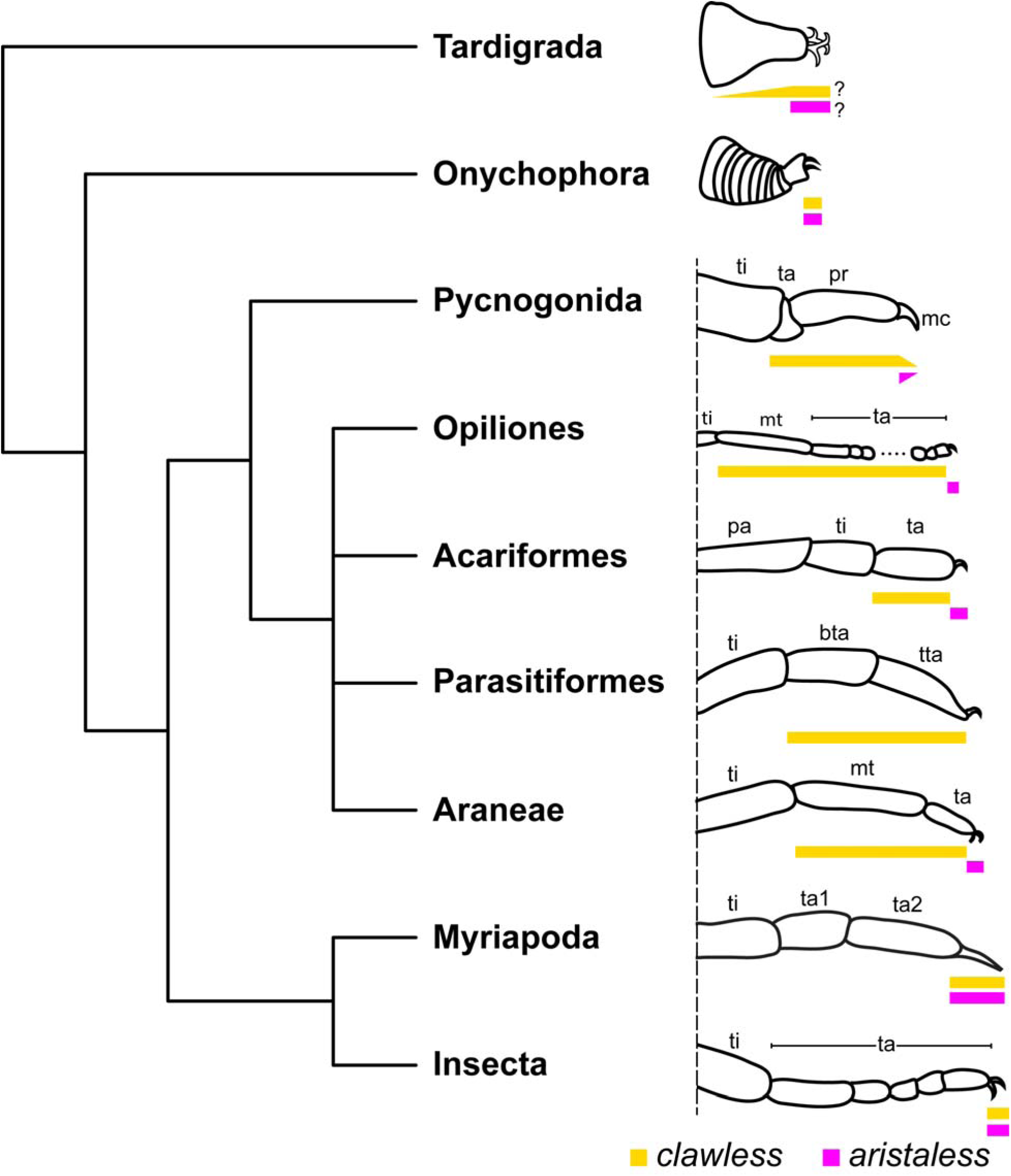
Developmental system drift in distal appendage patterning dynamics in Chelicerata. Schematic representations of distal podomere morphology and expression domains of cll and al in representative lineages of Panarthropoda. Note restricted distal domains of cll expression in Onychophora, Myriapoda, and Insecta. Proximal al domains have been excluded for simplicity. “?” refers to ambiguous expression domains from [40]. Abbreviations: bta – basitarsus; tta – telotarsus; others as in Figure 1.

The sum of these available datasets suggests that retention of cll expression in the distal tip of the developing appendages across Panarthropoda reflects an ancestral role in the patterning and formation of the claws. However, this function was lost in the chelicerate common ancestor following the split from Mandibulata, with no observable impact on the expression of the phenotype. This event is consistent with a case of developmental drift and highlights the lability of the gene regulatory network underlying the patterning of the proximo-distal appendage axis across panarthropods. Several cases of labile gene expression have been observed in the arthropod appendage literature, as exemplified by surveys of nubbin, homothorax, extradenticle, dachshund, Lim1, Sp6-9; and even Distal-less [4,22,41–48]. The variability in expression domains of these transcription factors even within a subset of arthropods (e.g., insects; arachnids) suggests that the patterning of the leg axis and its subdivisions is robust, possibly by way of redundant wiring of the underlying gene network [49]. Functional evidence for this redundancy is limited, though a recent example supporting its persistence is the redundancy of extradenticle and dachshund-2 function in patterning the distal segmental boundary of the patella in a harvestman and a spider, respectively [4,26].

More generally, this case study underscores the value of comparative data and enriching taxonomic sampling, even for clades with well-established model systems, toward closing gaps in the understanding of morphological evolution. With respect to chelicerates, two remaining taxon-specific innovations whose genetic underpinnings are still not understood include the three coxae of sea spiders (recently proposed to be subdivisions of the coxa of arachnids, based on the expression of SoxNeuro homologs) and the metatarsus (or sea spider “tarsus”), whose presence distinguishes the walking leg of most arachnids from the pedipalp.

## Authors’ Contributions

B.C.K.: conceptualization, data collection, data curation, formal analysis, writing – original draft, writing – review and editing. S.M.N.: data collection, formal analysis, writing – review and editing. E.M.L.: data collection, formal analysis, writing – review and editing. E.V.W.S.: data collection, formal analysis, writing – review and editing. I.A.H.: data collection, formal analysis, writing – review and editing. A.A.B.: data collection, formal analysis, writing – review and editing. M.H.: data collection, formal analysis, writing – review and editing. G.B.: conceptualization, data collection, formal analysis, funding acquisition, writing – original draft, writing – review and editing. M.G.-N.: funding acquisition, resources, supervision, writing – review and editing. P.P.S.: conceptualization, data collection, data curation, formal analysis, funding acquisition, project administration, writing – original draft, writing – review and editing.

## Acknowledgements

We are indebted to the Newcomb Imaging Center, Department of Botany, University of Wisconsin-Madison for imaging of P. opilio. Sarah Swanson assisted with confocal microscopy training and troubleshooting. Nithya Atla assisted with the care and maintenance of P. opilio. Tick imaging was performed at the High Spatial and Temporal Imaging Core (HSTRI, Core C), University of Nevada, Reno, which is supported by the National Institute of General Medical Services (P20GM130459). This work was supported by the National Science Foundation grant no. IOS-2016141 (to P.P.S.), the Austrian Science Fund (FWF) 10.55776/PAT1457924 (to G.B.), and the National Institutes of Health – National Institute of Allergy and Infectious Diseases (NIH-NIAID) USA grants R21AI176352 and R01AI172943 (to M.G.-N.).

---

**Figure S1.**
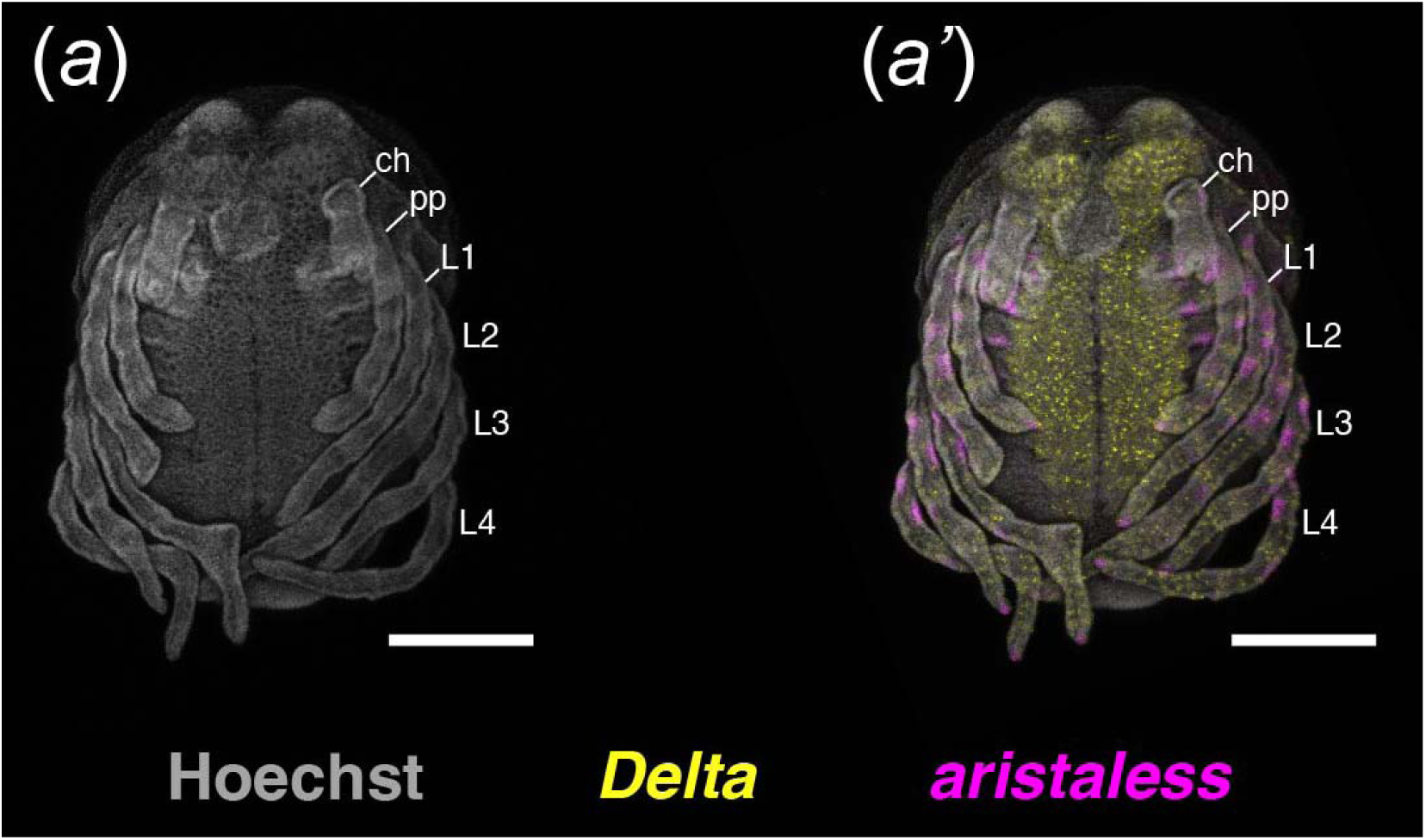
Severe Po-cll RNAi phenotypes are characterized by kinked appendages. (a) isolated nuclear counterstaining of stage 11 Po-cll RNAi embryo. (a’) Same embryo as in (a) with multiplexed expression of Po-Dl (yellow) and Po-al (magenta). Scale bars: 200 µm. Abbreviations as in figure 1.

**Figure S2.**
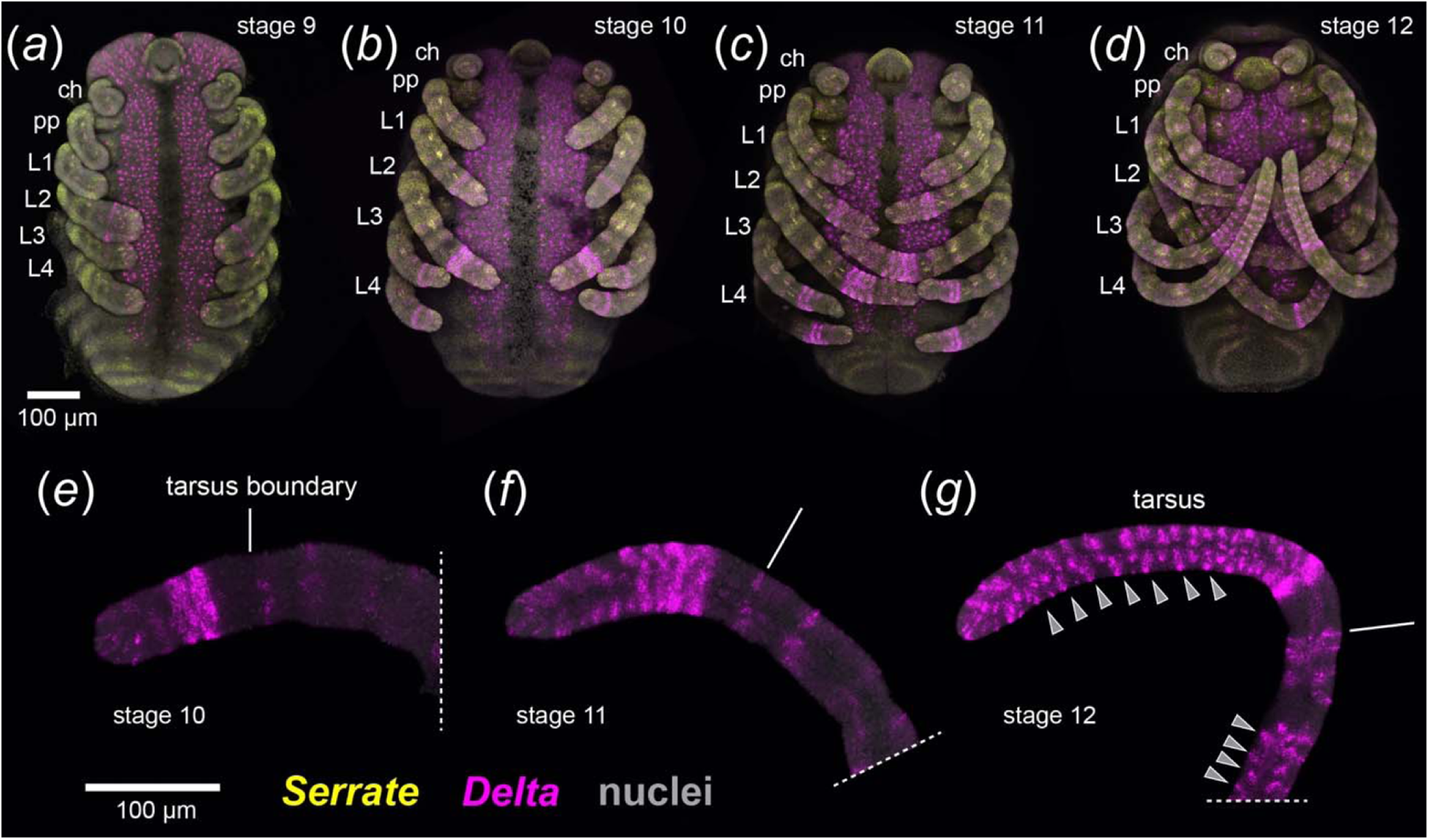
Characterization of Po-Dl and Po-Ser expression dynamics in the elongating tarsus of P. opilio. (a-d) Multiplexed expression of Po-Dl (magenta) and Po-Ser (yellow) with nuclear counterstaining (grey) in representative stages of tarsal elongation. (a) Stage 9 embryo. (b) Stage 10 embryo. (c) Stage 11 embryo. (d) Stage 12 embryo. (e-g) Leg II appendage mounts with isolated expression of Po-Dl. (e) Leg II of stage 10 embryo. (f) Leg II of stage 11 embryo. (g) Leg II of stage 12 embryo. Grey arrowheads: Po-Dl expression in putative sensory cells. Abbreviations as in figure 1.

**Figure S3.**
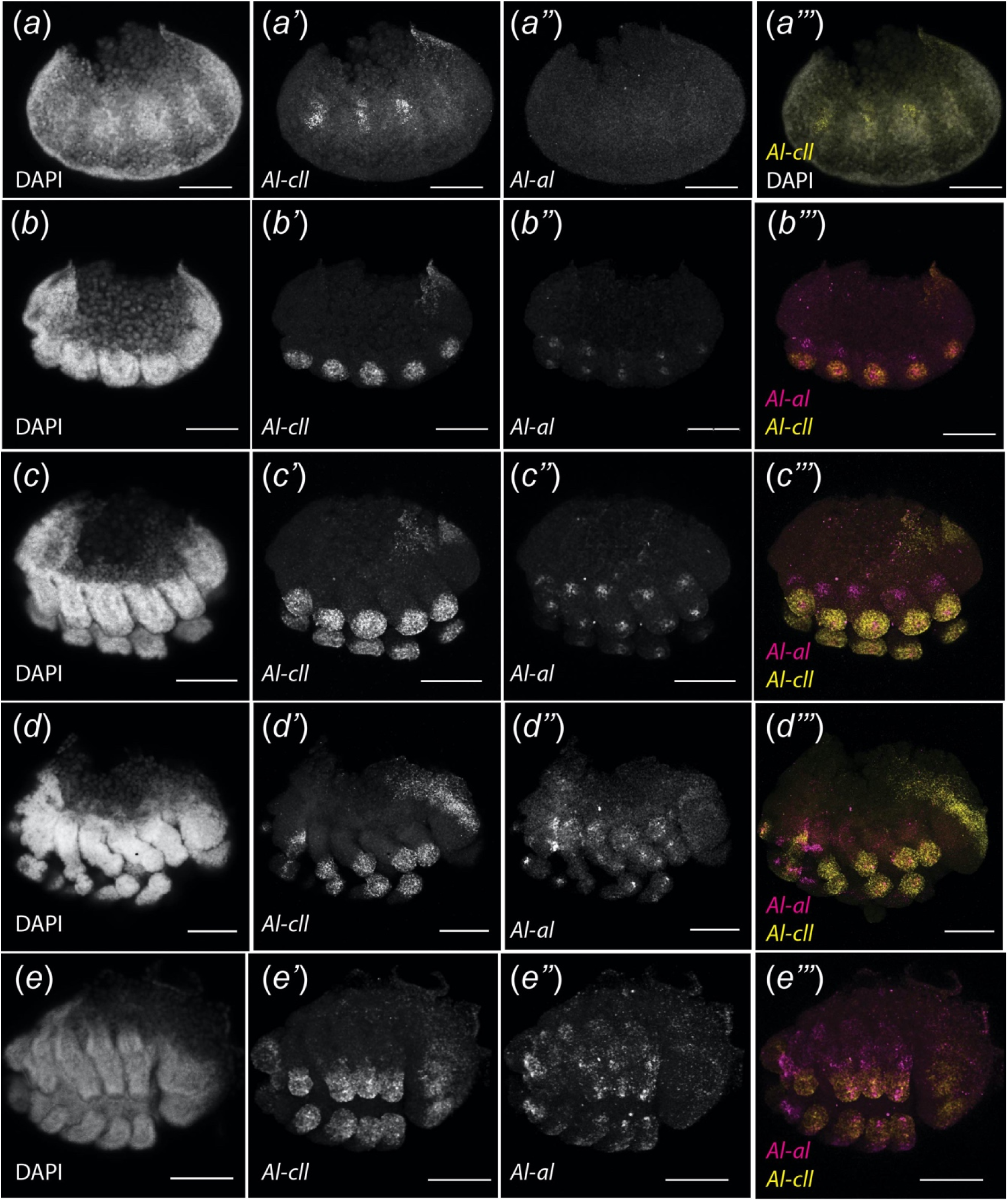
Expression dynamics of Al-cll and Al-al in representative stages of appendage formation and elongation in the acariform mite A. longisetosus. (a-e) Nuclear counterstaining of successive limb bud stages. (a’-e’) Isolated expression of Al-cll. (a’’-b’’) Isolated expression of Al-al. (a’’’-b’’’) Multiplexed expression of Al-cll (yellow) and Al-al (magenta). Scale bars: 50 µm.

**Figure S4.**
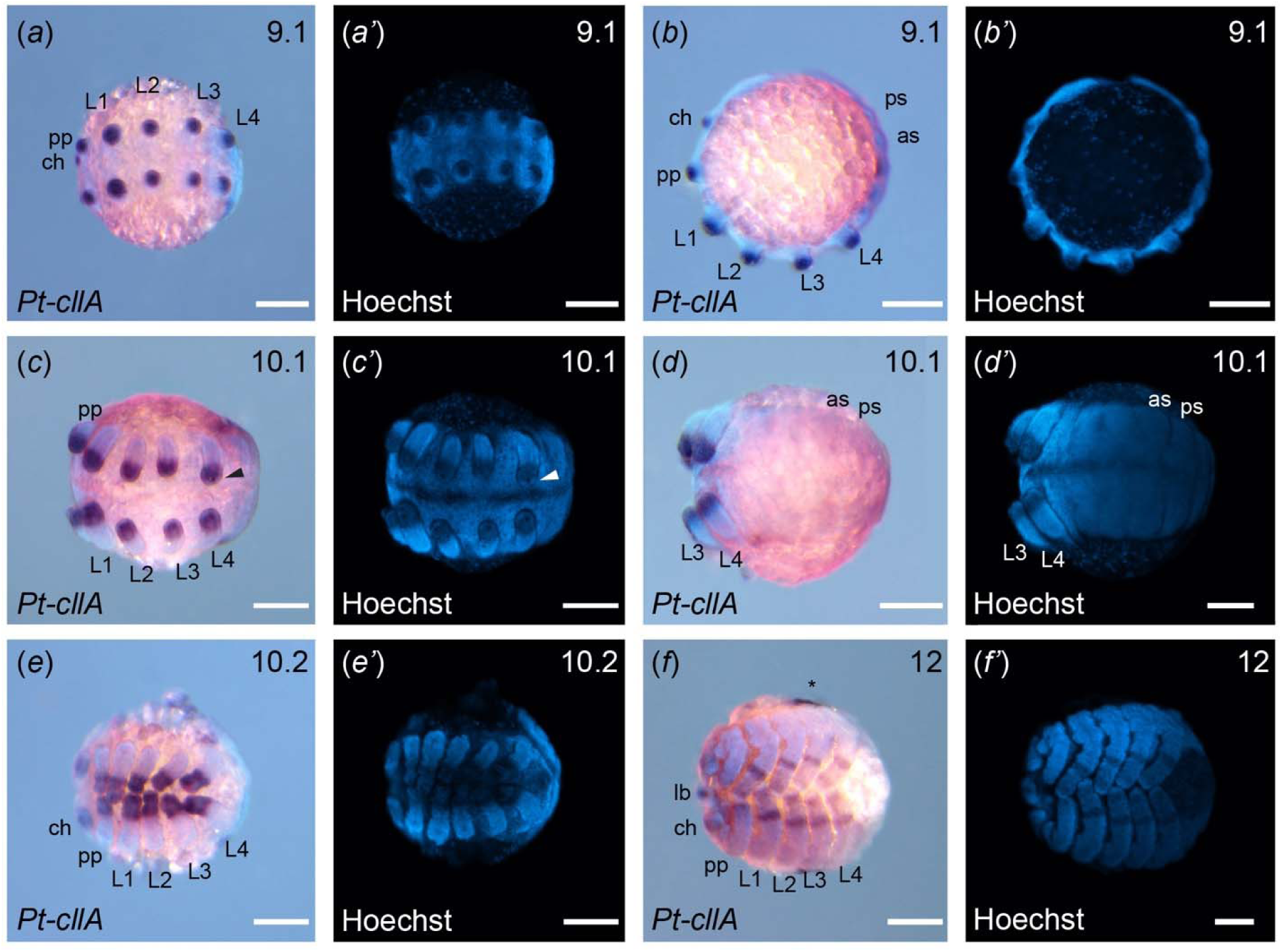
Pt-cllA expression dynamics during appendage formation and elongation in the spider P. tepidariorum. (a-f) Expression of Pt-cllA with nuclear counterstaining in P. tepidariorum embryos at stage 9.1 (a-b), 10.1 (c-d), 10.2 (e), and 12 (f). (a’-f’) Isolated nuclear counterstaining of same embryos depicted in (a-f). Scale bars: 100 µm. Abbreviations: as – anterior spinneret; ps – posterior spinneret; others as in figure 1. Asterisk: Non-specific staining.

**Figure S5.**
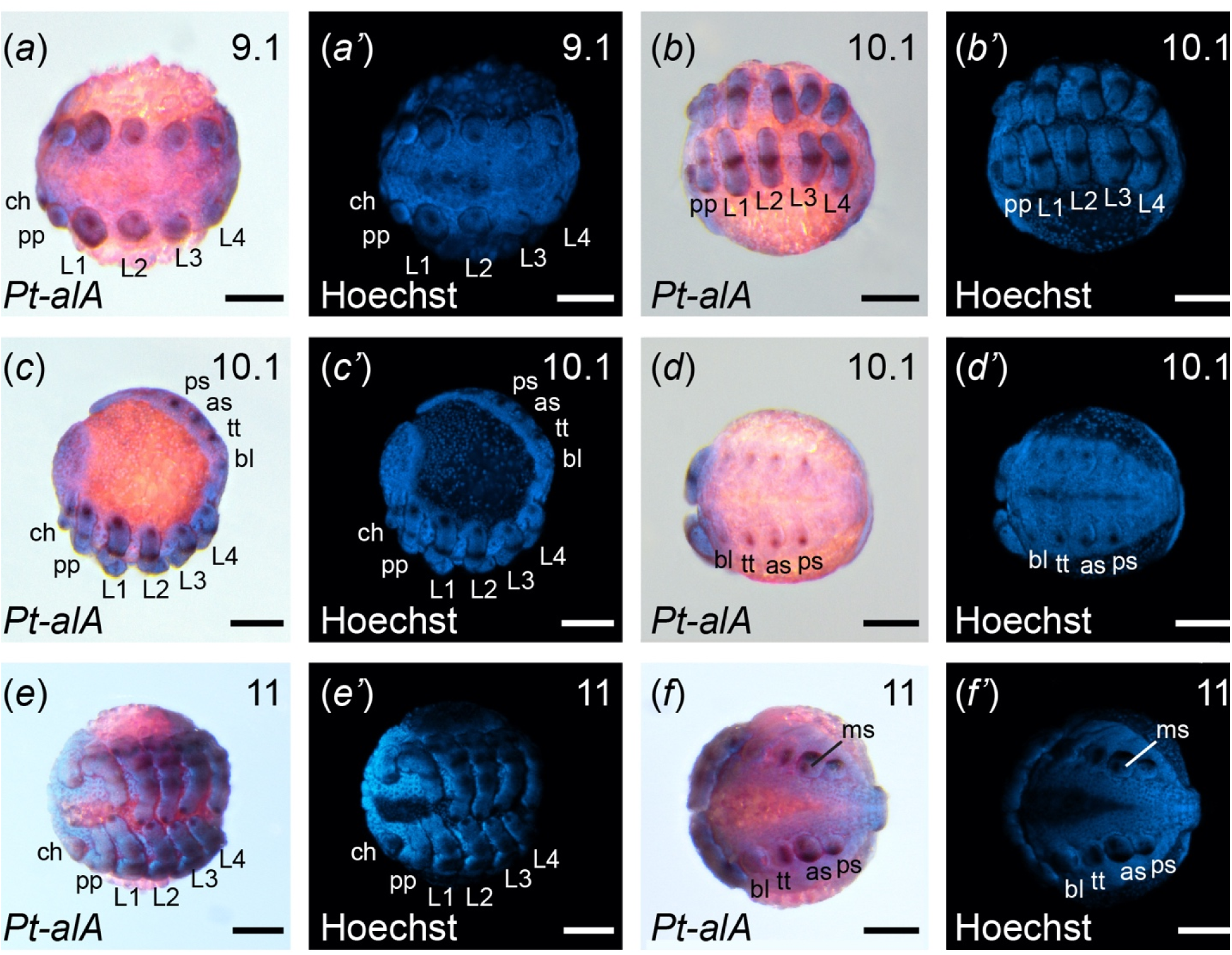
Pt-al expression dynamics during appendage formation and elongation in the spider P. tepidariorum. (a-f) Expression of Pt-al with nuclear counterstaining in P. tepidariorum embryos at stage 9.1 (a), 10.1 (b-d), 11 (e-f). (a’-f’) Isolated nuclear counterstaining of same embryos depicted in (a-f). Scale bars: 100 µm. Abbreviations: bl – book lung; tt – tubular tracheae; ms – median spinneret; others as in figure S4.

**Figure S6.**
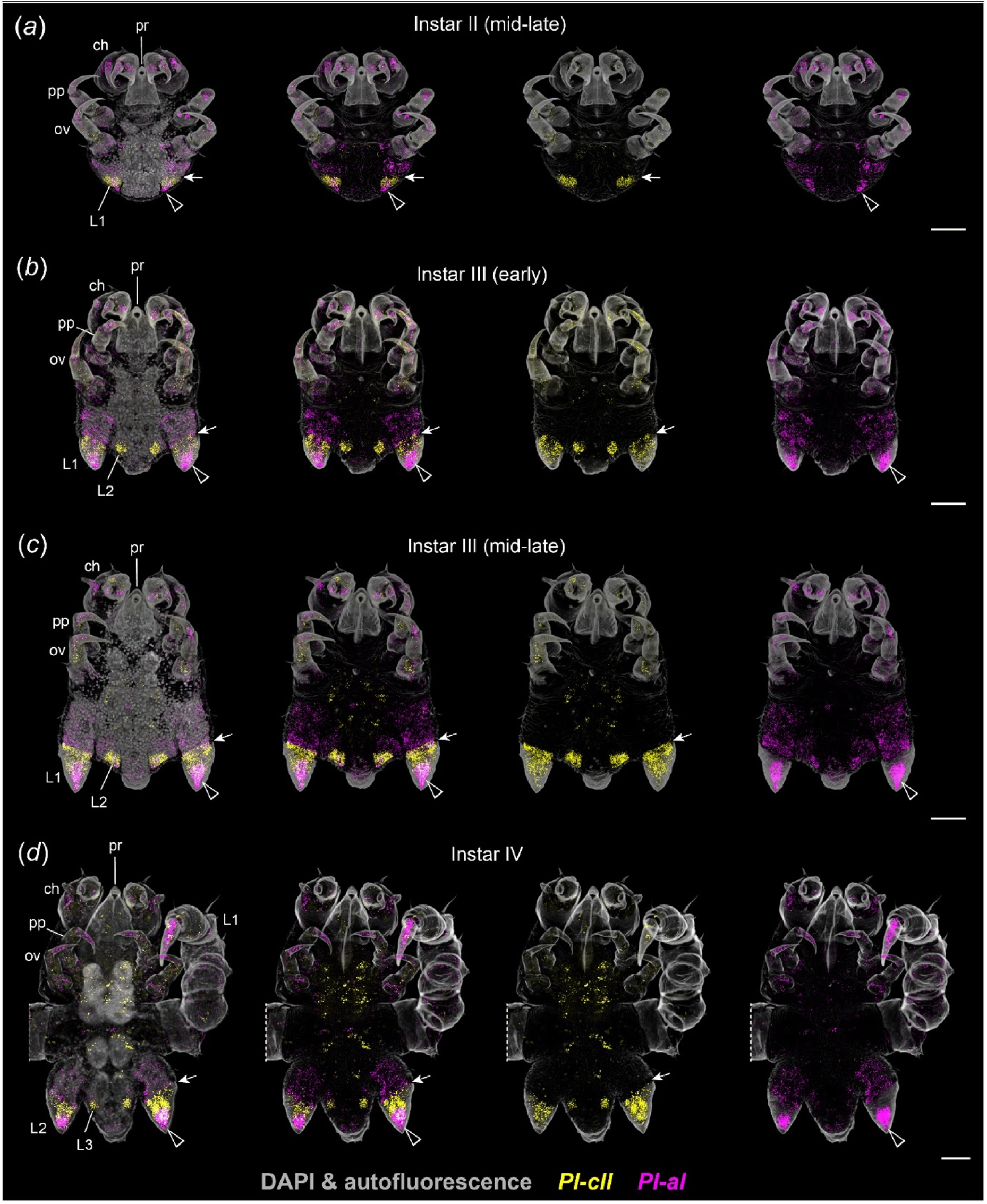
Pl-cll and Pl-al expression during postembryonic instars II-IV of the sea spider P. litorale. Maximum intensity projections of CLSM stacks in ventral view, cuticular autofluorescence and nuclear counterstaining in grey. (a) Instar II. (b) Early instar III. (c) Advanced instar III. (d) Instar IV. Scale bars: 50 µm. Abbreviations: ov – ovigeral larval limb; pr – proboscis; others as in figure S4. Black arrowhead: Pl-al positive domain in the distal tip territory of limb buds. Arrow: sharp proximal boundary of Pl-cll expression in limb buds.

**Figure S7.**
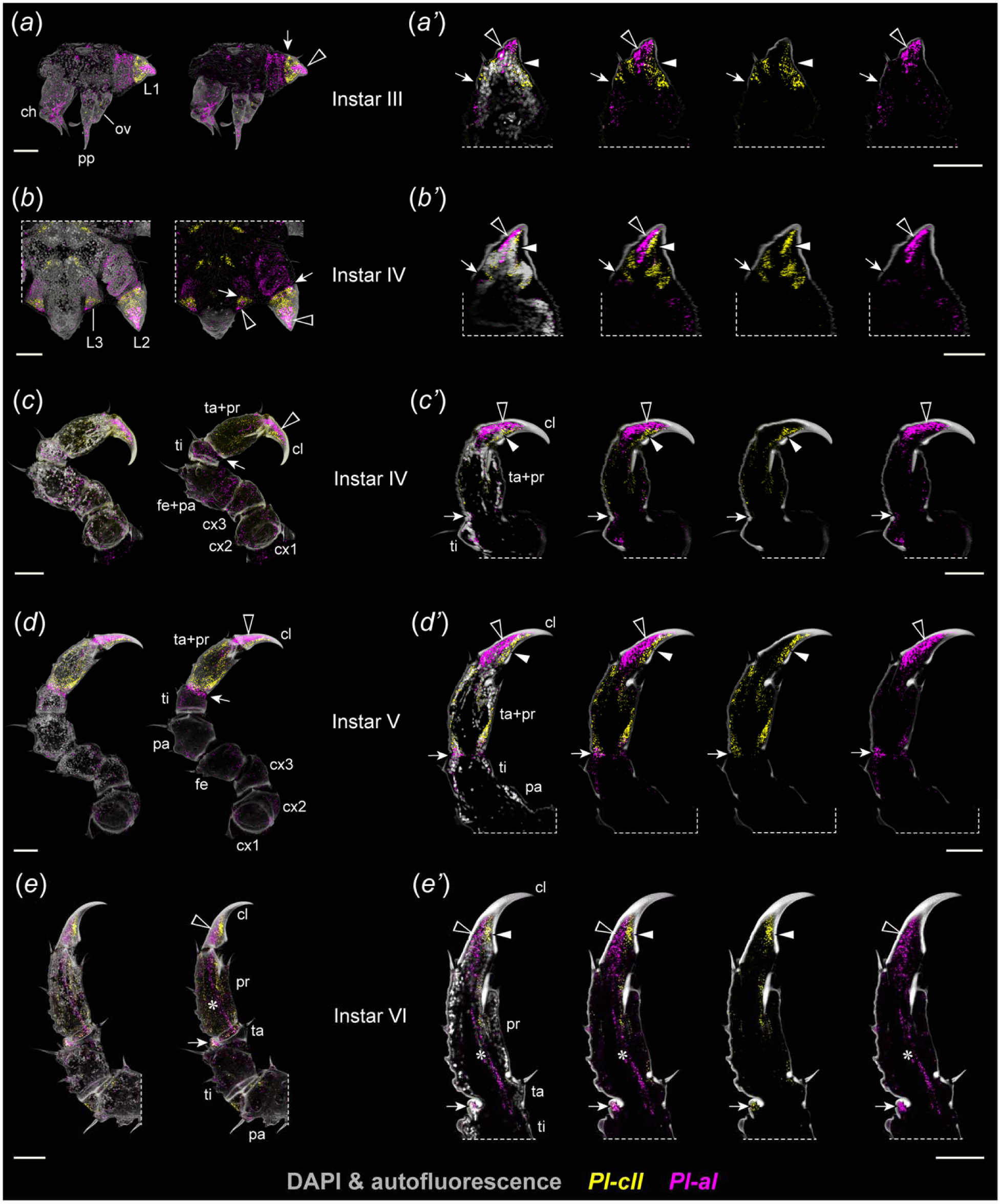
Details of Pl-cll and Pl-al expression dynamics during leg development of the sea spider P. litorale. Cuticular autofluorescence and nuclear counterstaining in grey. (a-e) Maximum intensity projections of complete CLSM stacks. (a’-e’) Extended virtual sections through limb buds and legs with ectal side pointing left. (a,a’) Instar III in lateral view (a) and leg 1 bud (a’). (b,b’) Instar IV. Ventral view of posterior body pole (b) and pre-molting leg 2 bud with internally folded leg tissue (b’). (c,c’) Instar IV. Leg 1 (c) and detail of its distal podomeres (c’). (d,d’) Advanced instar V. Leg 1 (d) and detail of its distal podomeres (d’). (e,e’) Instar VI, distal podomeres of leg 1. Scale bars: 50 µm. Abbreviations: ov – ovigeral limb bud; others as in figure 4. Black arrowhead: ectal Pl-al domain in the distal limb bud tips and in the main claw of legs. White arrowhead: mesal Pl-cll domain in the distal limb bud tips and in the main claw of legs. Arrow: proximal boundary of Pl-cll expression in limb buds and region of Pl-cll and Pl-al co-expression at the tibial-tarsal boundary in developing legs. Asterisk: Pl-al positive (putatively mesodermal) cell strand in the propodus of instar VI.

