## Supplementary File S5 for "Developmental system drift in the patterning of the arthropod tarsus"

**Table S1.** Probe pairs designed for *Phalangium opilio clawless* HCR *in situ* hybridization (B2 initiator).

| **Pair** | **Initiator** | **Spacer** | **Hybridization** | **Hybridization** | **Spacer** | **Initiator** |
| --- | --- | --- | --- | --- | --- | --- |
| 1 | CCTCGTAAATCCTCATCA | AA | ATGTACAAAATCGCGATACAACGCC | AAAAACTTAAACTAAGCGCTGGAAG | AA | ATCATCCAGTAAACCGCC |
| 2 | CCTCGTAAATCCTCATCA | AA | CGCAAAATGTTCCATAATGTTCAGC | TCCAAGCTGAATTTTTGTTGGTCAT | AA | ATCATCCAGTAAACCGCC |
| 3 | CCTCGTAAATCCTCATCA | AA | GCGCGTGTTCGAATTGTTTTAAACG | ATTAAAACACATGAAATTCTGGTAG | AA | ATCATCCAGTAAACCGCC |
| 4 | CCTCGTAAATCCTCATCA | AA | GTCGGTATTTTTAGTGTGGGAAAAA | AAAAAGGCGTGTTTAGCGTGTGACT | AA | ATCATCCAGTAAACCGCC |
| 5 | CCTCGTAAATCCTCATCA | AA | TCGGTTCTTTAGTACAATCGGAGGC | ATAATGATTTTTGTCCGTTTTCCGT | AA | ATCATCCAGTAAACCGCC |
| 6 | CCTCGTAAATCCTCATCA | AA | AATCGGAGGAATGTTTTAGTTTTAG | GTCCGAAAAATCGCGAAGCTCAATT | AA | ATCATCCAGTAAACCGCC |
| 7 | CCTCGTAAATCCTCATCA | AA | GAAGCATTTGTTTAGCCAAATTATC | AGGTGGTCATAATTCGTACTGAAAC | AA | ATCATCCAGTAAACCGCC |
| 8 | CCTCGTAAATCCTCATCA | AA | TCTGGCTAAGAGAGCGCGCATTTTA | ACCTCAACAGTGCTTCGATGACCTT | AA | ATCATCCAGTAAACCGCC |
| 9 | CCTCGTAAATCCTCATCA | AA | AATTGGGTGTGGGTTTATGCGAGTG | TAATTATAGTGGTAATTATTCCTTC | AA | ATCATCCAGTAAACCGCC |
| 10 | CCTCGTAAATCCTCATCA | AA | CAAATAGGTGGTGATGTGATATACG | TTCGAATCACTAGTTTAGGTTATTC | AA | ATCATCCAGTAAACCGCC |
| 11 | CCTCGTAAATCCTCATCA | AA | ATGATGGTGATGATGATGGCCAAGA | TGGTCGATCGGCCGAAGAGTGCACG | AA | ATCATCCAGTAAACCGCC |
| 12 | CCTCGTAAATCCTCATCA | AA | ACCGTTTGGTGGTCCGCTTCCCGGT | TCCGTTTTGACCATCTTGACCCATT | AA | ATCATCCAGTAAACCGCC |
| 13 | CCTCGTAAATCCTCATCA | AA | GGAGGGACGCGTTACTCAGACACAA | CCCAAGGTTGTAGGTTTTGTAGCGC | AA | ATCATCCAGTAAACCGCC |
| 14 | CCTCGTAAATCCTCATCA | AA | CCACTTCCGCTTGCAAGGACATCAT | TGGCCGACTCGTACAGGCTCTTGGA | AA | ATCATCCAGTAAACCGCC |
| 15 | CCTCGTAAATCCTCATCA | AA | GGCGGTTTGCCTTCTCCATTTAGTT | CGCTTGCCTTTCCGCTTCGCGTTCT | AA | ATCATCCAGTAAACCGCC |
| 16 | CCTCGTAAATCCTCATCA | AA | GCATCGGTCATTTTGAGTTGCTTGG | CTGTTCTGAAACCACGTCTTGACCT | AA | ATCATCCAGTAAACCGCC |
| 17 | CCTCGTAAATCCTCATCA | AA | ACTTTTGTTTGTGAAACCTCTTCTC | GAGCGGCCCTTTCGGCGGACGCCAA | AA | ATCATCCAGTAAACCGCC |
| 18 | CCTCGTAAATCCTCATCA | AA | CGTCCTGGGCTTTTTCCGTTTGGGA | TTCGCATATTTGCATTCTCGTAAAG | AA | ATCATCCAGTAAACCGCC |
| 19 | CCTCGTAAATCCTCATCA | AA | GCAAAGAGAATGGACCTGGTAGTCT | TCACCGGTGGTAACGCCGCTGATGA | AA | ATCATCCAGTAAACCGCC |
| 20 | CCTCGTAAATCCTCATCA | AA | TTCAGCTGTTGCGCGTGATGCTGAT | GCGCCTAAGACGGAGAACTTGAGTT | AA | ATCATCCAGTAAACCGCC |

**Table S2.** Probe pairs designed for *Phalangium opilio aristaless* HCR *in situ* hybridization (B1 initiator).

| **Pair** | **Initiator** | **Spacer** | **Hybridization** | **Hybridization** | **Spacer** | **Initiator** |
| --- | --- | --- | --- | --- | --- | --- |
| 1 | GAGGAGGGCAGCAAACGG | AA | TGGATTATATCCTTGATGGGCTTGG | TACTCCCGGTGGTACACCCGTTGAA | TA | GAAGAGTCTTCCTTTACG |
| 2 | GAGGAGGGCAGCAAACGG | AA | TTTGCACGACGATTTTGAAACCATA | CCAGCGGCCTTTTCTTGTTTACGCC | TA | GAAGAGTCTTCCTTTACG |
| 3 | GAGGAGGGCAGCAAACGG | AA | TCATAGCCAGTTCTTCTCTTGTGAA | GAACTCGTGCTTCGGTTAAATCGAC | TA | GAAGAGTCTTCCTTTACG |
| 4 | GAGGAGGGCAGCAAACGG | AA | TTTCTCAAGTTCTTCTAGTTGAAAA | GTCGGGATAGTGCGTCCTGCTGAAT | TA | GAAGAGTCTTCCTTTACG |
| 5 | GAGGAGGGCAGCAAACGG | AA | TTTCTTTTGGGAAAGTCCTCGGGTT | GTAAAGGTAGTTCTGTAACGTCTTT | TA | GAAGAGTCTTCCTTTACG |
| 6 | GAGGAGGGCAGCAAACGG | AA | GGTTTGGACGTCCCGAAGAGTTAAC | TTCCTCCACCGCCGCCGCCACCATT | TA | GAAGAGTCTTCCTTTACG |
| 7 | GAGGAGGGCAGCAAACGG | AA | CGGACTACCTGCATCTTGTTGTCTC | ATTGGCATCCCCGTTGTTGCTCAAC | TA | GAAGAGTCTTCCTTTACG |
| 8 | GAGGAGGGCAGCAAACGG | AA | TTAACCACGTCATCGCGTATCCCGT | CAATCCATCGGAGATTCCGACCGAT | TA | GAAGAGTCTTCCTTTACG |
| 9 | GAGGAGGGCAGCAAACGG | AA | TGTCCTCCGCGGAATTTCTTCGAGT | CAACCGAAAACTCTCCGTCGGAACA | TA | GAAGAGTCTTCCTTTACG |
| 10 | GAGGAGGGCAGCAAACGG | AA | GAATATTGTTAAGACCATGAACGCC | CGTGATGACTGTTATTGTTATTTTC | TA | GAAGAGTCTTCCTTTACG |
| 11 | GAGGAGGGCAGCAAACGG | AA | TAACGCGTTTGGACTTCCGGTTGTG | ACTTTTAAGTATTTTCTCCGCACTA | TA | GAAGAGTCTTCCTTTACG |
| 12 | GAGGAGGGCAGCAAACGG | AA | GCGCCAACGTGACGTAAGAATTGTT | GGACGCGGCGATGGTCCACCACCCG | TA | GAAGAGTCTTCCTTTACG |
| 13 | GAGGAGGGCAGCAAACGG | AA | GGAATAGTTCGAAGTTACGTTGATG | GCTGGTGCAAGTTTTGTAATTGTTG | TA | GAAGAGTCTTCCTTTACG |
| 14 | GAGGAGGGCAGCAAACGG | AA | ATGATGGTAATTGTTCCCGAGCATT | GGCCGCCGCCGCTAAATGGTGCGGG | TA | GAAGAGTCTTCCTTTACG |
| 15 | GAGGAGGGCAGCAAACGG | AA | CGTGCGGATTCGTGGGGATGATTAG | TCACTGGCTGATGTTAAGCCTAACG | TA | GAAGAGTCTTCCTTTACG |
| 16 | GAGGAGGGCAGCAAACGG | AA | TCCCCGTAACGGCAACGCGATTGTT | CCGATACACCCATGGTTGTTGTCGA | TA | GAAGAGTCTTCCTTTACG |

**Table S3.** Probe pairs designed for *Phalangium opilio Delta* HCR *in situ* hybridization (B2 initiator).

| **Pair** | **Initiator** | **Spacer** | **Hybridzation** | **Hybridzation2** | **Spacer3** | **Initiator4** |
| --- | --- | --- | --- | --- | --- | --- |
| 1 | CCTCGTAAATCCTCATCA | AA | CGCGCACGTATAAGATCCTTGTCCC | ATTGGTTCCCGAAAATCCATCAGGA | AA | ATCATCCAGTAAACCGCC |
| 2 | CCTCGTAAATCCTCATCA | AA | GGCTTGTGATTTGTGCAATAATTGA | TTCGTACAAGTACCGCCGTTCTTAC | AA | ATCATCCAGTAAACCGCC |
| 3 | CCTCGTAAATCCTCATCA | AA | CTTCGTCGCACGTACACTGCCAAGG | CTTGGTTACAAAAGAGACCACCCCA | AA | ATCATCCAGTAAACCGCC |
| 4 | CCTCGTAAATCCTCATCA | AA | ATAGCGAATACATTCGTCGCAACGA | GTTGCAAGTTCCGTGTTGACAACCA | AA | ATCATCCAGTAAACCGCC |
| 5 | CCTCGTAAATCCTCATCA | AA | TCGTTTGGCTTATTGCAGGTACCGT | CCTTGCCAACCTGTCCTACAAACGC | AA | ATCATCCAGTAAACCGCC |
| 6 | CCTCGTAAATCCTCATCA | AA | CTTTGGTACAATATTCGCCACTCCA | GTTGGTGGCAACCAGGAGCGCATAT | AA | ATCATCCAGTAAACCGCC |
| 7 | CCTCGTAAATCCTCATCA | AA | ACTACACGTATAATGTCCGAACTTG | AGGTAGACAGACTGTTTTACCCTCG | AA | ATCATCCAGTAAACCGCC |
| 8 | CCTCGTAAATCCTCATCA | AA | CCTTTACCGTAGTAATTAGGTTGGC | TCTCTGGGTCGACACAACTTGGTGC | AA | ATCATCCAGTAAACCGCC |
| 9 | CCTCGTAAATCCTCATCA | AA | CTTTACTGGTCTTGTTGTAATGTTC | GAACGCGGTAGGCGTACTTGACGGC | AA | ATCATCCAGTAAACCGCC |
| 10 | CCTCGTAAATCCTCATCA | AA | AGGTGTCGCCAAACGATAGATTAGA | CCAATTGGGACCGACGTCCAACCAA | AA | ATCATCCAGTAAACCGCC |
| 11 | CCTCGTAAATCCTCATCA | AA | CCATTTCTGGAGTTCCAAGCTTCGA | GCATTGGGACCAGGTGATGTAGAGT | AA | ATCATCCAGTAAACCGCC |
| 12 | CCTCGTAAATCCTCATCA | AA | ATTCGAACTCGAATTTCATGGGATT | TCAAGGAAAACGTACCCGTCCACGT | AA | ATCATCCAGTAAACCGCC |
| 13 | CCTCGTAAATCCTCATCA | AA | GTTAAATGGAAAGATTATGAGGCTA | GAACATTCCCGTCGAATCGGAGTTA | AA | ATCATCCAGTAAACCGCC |
| 14 | CCTCGTAAATCCTCATCA | AA | ACGCCATTAGGATCGATGGCGACTT | GGGGTAACTTTGGAACCGAATGTGC | AA | ATCATCCAGTAAACCGCC |
| 15 | CCTCGTAAATCCTCATCA | AA | TCCGGCAACTTCCGGTACACGATTT | AATGTTGAAGACATACTCGAAAGTA | AA | ATCATCCAGTAAACCGCC |
| 16 | CCTCGTAAATCCTCATCA | AA | TCGAAATTCGACATTCTCAAAAGAA | GTCGCAACAATCTCCTTGCGCGTCT | AA | ATCATCCAGTAAACCGCC |
| 17 | CCTCGTAAATCCTCATCA | AA | GAACATGACTGTGGCAATAGAGTTA | AAACGTAGCTCAAAAACACCTTCGC | AA | ATCATCCAGTAAACCGCC |
| 18 | CCTCGTAAATCCTCATCA | AA | ACGACGTTCGAGATCCAATGGCAAT | GCAACGTCGTTGTCAAAATAATAAC | AA | ATCATCCAGTAAACCGCC |

**Table S4.** Probe pairs designed for *Phalangium opilio Serrate* HCR *in situ* hybridization (B1 initiator).

| **Pair** | **Initiator** | **Spacer** | **Hybridzation** | **Hybridzation2** | **Spacer3** | **Initiator4** |
| --- | --- | --- | --- | --- | --- | --- |
| 1 | GAGGAGGGCAGCAAACGG | AA | TGTTTCAACTTTAACCTCGATAACG | AGTACTCGTTTCGGGACTGATTCGA | TA | GAAGAGTCTTCCTTTACG |
| 2 | GAGGAGGGCAGCAAACGG | AA | CTGCTGATTAAATCGGCCATTCTCT | GACAAGGCGACACTATCAGATAGTT | TA | GAAGAGTCTTCCTTTACG |
| 3 | GAGGAGGGCAGCAAACGG | AA | GTATGGAAGGTTCGGCTGTGGAAGA | CGGCTTCGCTTAAGACTTGAAAAGT | TA | GAAGAGTCTTCCTTTACG |
| 4 | GAGGAGGGCAGCAAACGG | AA | CCACATTGAACAATAATTGGGAACG | ACAGTATCATCTTCGTTCATTTTCA | TA | GAAGAGTCTTCCTTTACG |
| 5 | GAGGAGGGCAGCAAACGG | AA | GTCTAAGTTGGACGCAGACTTCTTC | GAGTGAAGTTGTGAAGTATTGGGAA | TA | GAAGAGTCTTCCTTTACG |
| 6 | GAGGAGGGCAGCAAACGG | AA | AGCGCAGTTATTGCTGAGAGTGGCG | ACTCTTATCAAATAATAGAGTAATT | TA | GAAGAGTCTTCCTTTACG |
| 7 | GAGGAGGGCAGCAAACGG | AA | TATGGTGAGGTTAACATTGGGTCGT | TTAGGTTCGCATTCGTTGTTGAGTT | TA | GAAGAGTCTTCCTTTACG |
| 8 | GAGGAGGGCAGCAAACGG | AA | CTCTGGGTTTACAAGGTGGCGTGAA | GGTCGTGTCCAATACGAGTACAAAT | TA | GAAGAGTCTTCCTTTACG |
| 9 | GAGGAGGGCAGCAAACGG | AA | CGAACAGATATGATTAGGTGGACAC | ATCAAGGGCACTTTGACGGTGAATA | TA | GAAGAGTCTTCCTTTACG |
| 10 | GAGGAGGGCAGCAAACGG | AA | GGAACACAGTTCGGATGTCCACACC | TGTACTCCGTCAATTGGGTTCTGAT | TA | GAAGAGTCTTCCTTTACG |
| 11 | GAGGAGGGCAGCAAACGG | AA | GCATCGACAAGTGTTACACTCCCCT | CTGAGTGCATGTCACGTGTCCACGG | TA | GAAGAGTCTTCCTTTACG |
| 12 | GAGGAGGGCAGCAAACGG | AA | TTTCCATCCCAGATACAAGAACCGG | ACCCAAGTACTGCCATCAGGTTGAG | TA | GAAGAGTCTTCCTTTACG |
| 13 | GAGGAGGGCAGCAAACGG | AA | ATCGGCCTCCTTTACCAGGTGGACA | TTACAATTGGTACAACATCACGACA | TA | GAAGAGTCTTCCTTTACG |
| 14 | GAGGAGGGCAGCAAACGG | AA | GCAGGTACTGCCATCGGCGCAGGGA | ACAAGTGACGTTATTGATCTCATCT | TA | GAAGAGTCTTCCTTTACG |
| 15 | GAGGAGGGCAGCAAACGG | AA | CATTTGGGTCCAGTAAAACCAGGCG | CTATTACATTCGTTTTTATTGATCC | TA | GAAGAGTCTTCCTTTACG |
| 16 | GAGGAGGGCAGCAAACGG | AA | CAATACAACGACCACCGTTCATACA | ATTCGCACAAGTACCAATTAACTCC | TA | GAAGAGTCTTCCTTTACG |
| 17 | GAGGAGGGCAGCAAACGG | AA | TTGACACTTATGTCCATCATAACCT | ATTAGGTGAGCAGTCGTCAACGTTG | TA | GAAGAGTCTTCCTTTACG |
| 18 | GAGGAGGGCAGCAAACGG | AA | ACCGTGACGCATGTTGCGCCATTTT | CGACAGACGCAACTGTACGAATTAC | TA | GAAGAGTCTTCCTTTACG |
| 19 | GAGGAGGGCAGCAAACGG | AA | TAGAACTCTGACATGTCTTTCCTTG | AAGGATTTGATCTGCAGCTTGGATT | TA | GAAGAGTCTTCCTTTACG |
| 20 | GAGGAGGGCAGCAAACGG | AA | GTCACCGATATCGACGCAACTACCG | ACCTTCTTTGCAATGACACACAAAA | TA | GAAGAGTCTTCCTTTACG |
| 21 | GAGGAGGGCAGCAAACGG | AA | TGTGTCTTAAATTGCAAATTTTTCC | TTTGACACGTCGTGATATCGCAATG | TA | GAAGAGTCTTCCTTTACG |
| 22 | GAGGAGGGCAGCAAACGG | AA | ATCATTGATTAAATCGTTGCAAACA | CCATGGATGTTTGCACTCGCATGTA | TA | GAAGAGTCTTCCTTTACG |
| 23 | GAGGAGGGCAGCAAACGG | AA | TCGTTCTTATTTATATTGCAAACGG | TCATGTCGACATGGATTCGGATCGC | TA | GAAGAGTCTTCCTTTACG |
| 24 | GAGGAGGGCAGCAAACGG | AA | GGAATGTATTAACACCATCGACGCA | CTTCCCATCCATCAGGACAAACGCA | TA | GAAGAGTCTTCCTTTACG |
| 25 | GAGGAGGGCAGCAAACGG | AA | GCAATCATTTATATTCTCGTGACAG | CCCACCGTTTTGACAAGGATTCGAT | TA | GAAGAGTCTTCCTTTACG |
| 26 | GAGGAGGGCAGCAAACGG | AA | AAGTAAAACCACCATTTGGTTCGCT | TGCCGCTGTACCCTGGATCGCAGAC | TA | GAAGAGTCTTCCTTTACG |
| 27 | GAGGAGGGCAGCAAACGG | AA | AGTTGTTACCATGTGGTATCTTGAA | GCATTTTCCGTGGTTACCGCAAACT | TA | GAAGAGTCTTCCTTTACG |
| 28 | GAGGAGGGCAGCAAACGG | AA | TCAGTTTCTGAACAAGGTGGATTGT | CCTGAAGAGACTGGTACCATACAAC | TA | GAAGAGTCTTCCTTTACG |

**Table S5.** Probe pairs designed for *Archegozetes longisetosus clawless* HCR *in situ* hybridization (B2 initiator).

| **Pair** | **Initiator** | **Spacer** | **Hybridization** | **Hybridization** | **Spacer** | **Initiator** |
| --- | --- | --- | --- | --- | --- | --- |
| 1 | CCTCGTAAATCCTCATCA | AA | GTAAGTTGTACTGTACAGAGATTTC | AAATGTTTGATAGCGATCGATGCAA | AA | ATCATCCAGTAAACCGCC |
| 2 | CCTCGTAAATCCTCATCA | AA | TGATGCCAACGGAAACGGTGGTCTA | ATTCGAGTCTAGTAACTAACAGATT | AA | ATCATCCAGTAAACCGCC |
| 3 | CCTCGTAAATCCTCATCA | AA | CCGTAGATCGATTTGGATACAACTT | GGGTGGTCAGATCCGGGCTGACCAG | AA | ATCATCCAGTAAACCGCC |
| 4 | CCTCGTAAATCCTCATCA | AA | CTGCCTGACGTTCAGCCTCTCTTTC | CCTGTAATGACATCATCAGTCTATT | AA | ATCATCCAGTAAACCGCC |
| 5 | CCTCGTAAATCCTCATCA | AA | CCTATTTTGAAACCAAGTCTTGACT | TGCTGTCTGACGTCTCCACTTTGTT | AA | ATCATCCAGTAAACCGCC |
| 6 | CCTCGTAAATCCTCATCA | AA | AGTGATGCTCTTTCAGCAGATGCTA | GCGTCTGTCATTTTCAATTGTTTGG | AA | ATCATCCAGTAAACCGCC |
| 7 | CCTCGTAAATCCTCATCA | AA | TTGCATTCTTGTGAATGATGTGCGC | GTGAAATCTCTTTTCTAGTTCACAA | AA | ATCATCCAGTAAACCGCC |
| 8 | CCTCGTAAATCCTCATCA | AA | TGATAGGGATGACCTATTCTTCGAG | TTCTTCCTCTTTGGTGGTGTACGGT | AA | ATCATCCAGTAAACCGCC |
| 9 | CCTCGTAAATCCTCATCA | AA | GAAAGGGCATTTGTAAACCGTCTTT | GTACTGGCGGTAGACCAGCGCCAAG | AA | ATCATCCAGTAAACCGCC |
| 10 | CCTCGTAAATCCTCATCA | AA | TGGGTAATGTGTAGTGTAAGGCATA | AAGCGTTGGCGGAGTTGGTCCTAAC | AA | ATCATCCAGTAAACCGCC |
| 11 | CCTCGTAAATCCTCATCA | AA | CCGGTCAGATGATTCAATGCTGCCT | ACACGTATGACCGAAGCGGCACCTG | AA | ATCATCCAGTAAACCGCC |
| 12 | CCTCGTAAATCCTCATCA | AA | TCAGATTACATTTAGCATCAGGTGA | ATGGACTCACTGTCACACGGCTTGT | AA | ATCATCCAGTAAACCGCC |
| 13 | CCTCGTAAATCCTCATCA | AA | ATTGTTTTCAGTTGATGGACATTGA | CGAAGAGGGGTATGAGGCAGCGGTC | AA | ATCATCCAGTAAACCGCC |
| 14 | CCTCGTAAATCCTCATCA | AA | CCTTCTGAAGTAGCGCCGCTGTGAT | CAAAGCGATGACGGAGACTCTGATT | AA | ATCATCCAGTAAACCGCC |
| 15 | CCTCGTAAATCCTCATCA | AA | ATAGTCGACTTATACTGAATGACAG | GTTTGTTTGAGCCACTATTTGCACC | AA | ATCATCCAGTAAACCGCC |
| 16 | CCTCGTAAATCCTCATCA | AA | GTTTTGATGCACTGGTGATGAGAGT | CGATGGTTTGCTAGAATTGATTGAA | AA | ATCATCCAGTAAACCGCC |
| 17 | CCTCGTAAATCCTCATCA | AA | TCCTCATCATCGTTAATGTCAATAC | TCTCGCTTTGGCGATCGAGACATAT | AA | ATCATCCAGTAAACCGCC |
| 18 | CCTCGTAAATCCTCATCA | AA | CTGATCGCGGCATTGAGTCCTCGTA | CATCTGAGTTTATGGGACTCGAAGA | AA | ATCATCCAGTAAACCGCC |
| 19 | CCTCGTAAATCCTCATCA | AA | TGTTGTGATGTTCACTTACAGGTAC | GATGAGAACAATTAAAAGCACATTT | AA | ATCATCCAGTAAACCGCC |
| 20 | CCTCGTAAATCCTCATCA | AA | TTGTTGTCTGCGTAAAAGTTAACGC | TTTGAGAAACAACTCCAAAACAAAC | AA | ATCATCCAGTAAACCGCC |

**Table S6.** Probe pairs designed for *Archegozetes longisetosus* *aristaless* HCR *in situ* hybridization (B3 initiator).

| **Pair** | **Initiator** | **Spacer** | **Hybridization** | **Hybridization** | **Spacer** | **Initiator** |
| --- | --- | --- | --- | --- | --- | --- |
| 1 | GTCCCTGCCTCTATATCT | TT | ATTGATGGGATCACAGAAAAAGTGT | TTTTTGTTGCCATTAATCAATGGAC | TT | CCACTCAACTTTAACCCG |
| 2 | GTCCCTGCCTCTATATCT | TT | TCGTGTTCGCGTGCTTTCAGTCGGA | GGGTCTTTGCTTCGCATAACTGCCA | TT | CCACTCAACTTTAACCCG |
| 3 | GTCCCTGCCTCTATATCT | TT | AGTCGCAGAGGAGGTGTCTGTCTGT | AGACCTTCGTTTGGACTCGAGTTCG | TT | CCACTCAACTTTAACCCG |
| 4 | GTCCCTGCCTCTATATCT | TT | AGAGAGAGATGCGAGAAGACTTTGG | TCCTATTTTGGGCCTATTATGTGAT | TT | CCACTCAACTTTAACCCG |
| 5 | GTCCCTGCCTCTATATCT | TT | GTTGAGGATATGCTGCCGTCAGTTC | TGTTTGCGGACGGATTGCCGTGAAA | TT | CCACTCAACTTTAACCCG |
| 6 | GTCCCTGCCTCTATATCT | TT | TTAAGGCTTGGATGATCTAATAGTG | GCTGGAGTATTCAAGTAACCCGCTG | TT | CCACTCAACTTTAACCCG |
| 7 | GTCCCTGCCTCTATATCT | TT | GTTGCACCACTCGGTGTTCCGCCAC | TTGGTGATCACTTGTGATGGCAACA | TT | CCACTCAACTTTAACCCG |
| 8 | GTCCCTGCCTCTATATCT | TT | TCGCAAGTTCCTCCCTCGTAAATAC | CTCGTGCTTCAGTTAAATCGACTCG | TT | CCACTCAACTTTAACCCG |
| 9 | GTCCCTGCCTCTATATCT | TT | CGTTTGGGGAAATCATCGGAATGAT | AAAGTCGTTCGATATCTTCGCTGCT | TT | CCACTCAACTTTAACCCG |
| 10 | GTCCCTGCCTCTATATCT | TT | CTAACTCGTCAGGACCGCTACGGGG | CACCACTGTTTCTAATGGTCTGTGA | TT | CCACTCAACTTTAACCCG |
| 11 | GTCCCTGCCTCTATATCT | TT | ATCGCGCTCGTCATCATCTACTTTG | CAAAGGATTATCGTCATCATCCGAA | TT | CCACTCAACTTTAACCCG |
| 12 | GTCCCTGCCTCTATATCT | TT | GACCGGGGATACAACGCCCTTATTA | GTCGTCATCAACATCGGATCTGCTC | TT | CCACTCAACTTTAACCCG |
| 13 | GTCCCTGCCTCTATATCT | TT | CACATCTTTCAAGTACTCCCATTCC | TTTTGTCACTTAATAAGTTTTTCGG | TT | CCACTCAACTTTAACCCG |
| 14 | GTCCCTGCCTCTATATCT | TT | GTTGTTATTGAGCCAATTGTTGGCG | GATCTCTTAATAGCCTTATTATTAT | TT | CCACTCAACTTTAACCCG |
| 15 | GTCCCTGCCTCTATATCT | TT | AAGACAGCGCTTCCGATTGTGTTGG | TGCAGCTGTGGGTCGGTTTGTTGAT | TT | CCACTCAACTTTAACCCG |
| 16 | GTCCCTGCCTCTATATCT | TT | TCGCATGAAATCCAGGCCTGTTGTA | TGGTGATGGTGTGGCCAAAGTGGTG | TT | CCACTCAACTTTAACCCG |
| 17 | GTCCCTGCCTCTATATCT | TT | GGCTTCTGGTGGTATGTAGTCGTTG | GACGGCGAAATGGACGGAATTATAT | TT | CCACTCAACTTTAACCCG |
| 18 | GTCCCTGCCTCTATATCT | TT | TTGACTCCTTTTCCAAACCAATTAT | TCGTGATTTCTGTCAAAGGAGCCAG | TT | CCACTCAACTTTAACCCG |
| 19 | GTCCCTGCCTCTATATCT | TT | AAGTTTTGGCAATTGGGTCTGAACC | CAGTGAGAAGTTGTTTGTTTTAAAG | TT | CCACTCAACTTTAACCCG |
| 20 | GTCCCTGCCTCTATATCT | TT | GTGAGGCACTCACACAGAAAGTCAT | TTTTTAATAAGAGAAGAACAGTTTG | TT | CCACTCAACTTTAACCCG |

**Table S7.** Probe pairs designed for *Ixodes scapularis clawless* HCR *in situ* hybridization (B1 initiator).

| **Pair** | **Initiator** | **Spacer** | **Hybridization** | **Hybridization** | **Spacer** | **Initiator** |
| --- | --- | --- | --- | --- | --- | --- |
| 1 | GAGGAGGGCAGCAAACGG | AA | TCCCTGGCGCCCTCGTAGATGGACT | AGCGAGGCGTTGCTCATGCAGAGCG | TA | GAAGAGTCTTCCTTTACG |
| 2 | GAGGAGGGCAGCAAACGG | AA | TGAGTCTGTTGGCGGCGTGGCGTTC | TCACCACCTCAGCCTGGAGTGACAT | TA | GAAGAGTCTTCCTTTACG |
| 3 | GAGGAGGGCAGCAAACGG | AA | ATCGTCAGGAAATGCCGAGTACGTT | TTCGCGCTCTTCGGCCGTCTGTCTC | TA | GAAGAGTCTTCCTTTACG |
| 4 | GAGGAGGGCAGCAAACGG | AA | CTGCGGTTCTGGAACCACGTCTTGA | AGAGGGCTGATTTTTCTCCACTTTG | TA | GAAGAGTCTTCCTTTACG |
| 5 | GAGGAGGGCAGCAAACGG | AA | TGCTTGTGGAAGCGCTTCTCGAGCT | GCCCGCTCCGCACTGGCCAGGTACT | TA | GAAGAGTCTTCCTTTACG |
| 6 | GAGGAGGGCAGCAAACGG | AA | AAGGTGGAGCGGGCTGAGCTGTGCT | CACGCGAATCACAGTCCCGTTGGCA | TA | GAAGAGTCTTCCTTTACG |

**Table S8.** Probe pairs designed for *Pycnogonum litorale clawless* HCR *in situ* hybridization (B5 initiator).

| **Pair** | **Initiator** | **Spacer** | **Hybridization** | **Hybridization** | **Spacer** | **Initiator** |
| --- | --- | --- | --- | --- | --- | --- |
| 1 | CTCACTCCCAATCTCTAT | AA | CGATTCACCTGTAATGCTAACCGTC | GGAATCATACGCTCGATGATGCTAT | AA | CTACCCTACAAATCCAAT |
| 2 | CTCACTCCCAATCTCTAT | AA | AGGTAAATTACTGGCCTCAGTTAAA | TGCGGTGAGCACCACTGGAACGATT | AA | CTACCCTACAAATCCAAT |
| 3 | CTCACTCCCAATCTCTAT | AA | TATTGGAGAGGCAAGCGTTGGTACA | CTAAGATAAATCTGAAGTCGATCAG | AA | CTACCCTACAAATCCAAT |
| 4 | CTCACTCCCAATCTCTAT | AA | GCGTTGTCGTGATTATTTTCTTGAC | CTCAGATAAGACGATGTGGAATCTC | AA | CTACCCTACAAATCCAAT |
| 5 | CTCACTCCCAATCTCTAT | AA | TGTTAATACATAATGGATCACCTCG | TGCTTTGCAAAGCGTGTAATGAAGC | AA | CTACCCTACAAATCCAAT |
| 6 | CTCACTCCCAATCTCTAT | AA | TACAGCTTCTGCTTGTAATGACATC | TGCAGGATCGCCATAGATTCCCTTG | AA | CTACCCTACAAATCCAAT |
| 7 | CTCACTCCCAATCTCTAT | AA | TCACGCTCTTCTGCTGTCTGTCTTC | AAACGATTAGCCGCTTGACGCTCTG | AA | CTACCCTACAAATCCAAT |
| 8 | CTCACTCCCAATCTCTAT | AA | CTTTACTTGCGCATCGGTCATCTTC | TTTAGTTCTCCTGTTTTGAAACCAC | AA | CTACCCTACAAATCCAAT |
| 9 | CTCACTCCCAATCTCTAT | AA | TTATGGAATCTCTTTTCCAATTCGC | CTTTCAGCGGACGCTAAATATTTCT | AA | CTACCCTACAAATCCAAT |
| 10 | CTCACTCCCAATCTCTAT | AA | GCTTCTTACGTTTCGGAGGTGTTCT | TCTGCATCCGAGTGAAAGATGTCCT | AA | CTACCCTACAAATCCAAT |
| 11 | CTCACTCCCAATCTCTAT | AA | CATCGGTACCGGAACTTGCATTCTG | CTGATACGGGTGACCTATTCTTCTA | AA | CTACCCTACAAATCCAAT |
| 12 | CTCACTCCCAATCTCTAT | AA | GCCATCCAAGGAAACGCAGCTGAAG | TTCATTGTGAGAGCTAAACTTCTTG | AA | CTACCCTACAAATCCAAT |
| 13 | CTCACTCCCAATCTCTAT | AA | TGTGCCGGTACCCTGATAACAGTTC | GAAAACTGCGGCATTCCGATCGGGT | AA | CTACCCTACAAATCCAAT |
| 14 | CTCACTCCCAATCTCTAT | AA | GTGATAAAGAATGCGACGTTATACC | GGGCACCACCAAAAAGTGGATCGTA | AA | CTACCCTACAAATCCAAT |
| 15 | CTCACTCCCAATCTCTAT | AA | ATCGGCGGTGATGCCATCGAACTGT | AGACTAGTAACCGGTATGTAAGCTC | AA | CTACCCTACAAATCCAAT |
| 16 | CTCACTCCCAATCTCTAT | AA | TACCGTTGCTACTTGCCGGTGACGA | TAAACGCCATGAAATGTTCAATCTG | AA | CTACCCTACAAATCCAAT |
| 17 | CTCACTCCCAATCTCTAT | AA | TTCGCCCAGTAGACGACTGATACTG | TGCAGAGTTGTCATCTTTATCTTCT | AA | CTACCCTACAAATCCAAT |
| 18 | CTCACTCCCAATCTCTAT | AA | GATGAAGATCTGTTTATTGTTGGAG | TTATTATTTGATGACGGTGAACGAG | AA | CTACCCTACAAATCCAAT |
| 19 | CTCACTCCCAATCTCTAT | AA | TGATTTCCATATTATCCGGATCCAT | GGGGAGTTTTCTCCGACCCAACATC | AA | CTACCCTACAAATCCAAT |

**Table S9.** Probe pairs designed for *Pycnogonum litorale aristaless* HCR *in situ* hybridization (B2 initiator).

| **Pair** | **Initiator** | **Spacer** | **Hybridization** | **Hybridization** | **Spacer** | **Initiator** |
| --- | --- | --- | --- | --- | --- | --- |
| 1 | CCTCGTAAATCCTCATCA | AA | ATGTTACTGATTGTCTCATTGGCTC | TTTTTCAAAGTACGGTTCTCACTGG | AA | ATCATCCAGTAAACCGCC |
| 2 | CCTCGTAAATCCTCATCA | AA | TCCGCCCATACCTGACGTAGCGATA | ACTTCTGAGTTTTTCGGCATCCGCG | AA | ATCATCCAGTAAACCGCC |
| 3 | CCTCGTAAATCCTCATCA | AA | TCGCCATCGCTTGATGACGACTCTG | TGCTCGCACTTCCGGTATTCGCTGA | AA | ATCATCCAGTAAACCGCC |
| 4 | CCTCGTAAATCCTCATCA | AA | CGCCAACATTTCGGCCAACAGAATC | TTCGAGTTGTTGCATCGCCTCGCGT | AA | ATCATCCAGTAAACCGCC |
| 5 | CCTCGTAAATCCTCATCA | AA | CAACGAGGCACTCGGTCCAGGAAAT | CATCGAAGCGGCCGAATTGATGTGC | AA | ATCATCCAGTAAACCGCC |
| 6 | CCTCGTAAATCCTCATCA | AA | GTAGACCGGTCCATTGACGGAATCT | ATAACGAACCGGCCGTCATGGTGGA | AA | ATCATCCAGTAAACCGCC |
| 7 | CCTCGTAAATCCTCATCA | AA | AGTAAAGGCCATCGAGGCGGATCGT | CAGTTCTCATGAATGGATTGTACAT | AA | ATCATCCAGTAAACCGCC |
| 8 | CCTCGTAAATCCTCATCA | AA | TCGGTGGTACAGACTTACATTCTCG | AACTTGTGGAACGTGAGGAACTCTC | AA | ATCATCCAGTAAACCGCC |
| 9 | CCTCGTAAATCCTCATCA | AA | TTGATGCGCCGGAATCGGTCCTTGA | CTTGTCTAAAGACGGAAGCCATTTC | AA | ATCATCCAGTAAACCGCC |
| 10 | CCTCGTAAATCCTCATCA | AA | GGATGTTCGTGCGGACCGACTTTCT | GACGCGTTCGATGGAAATCCGGAAT | AA | ATCATCCAGTAAACCGCC |
| 11 | CCTCGTAAATCCTCATCA | AA | GGAACCAAACCTGCACTCTAGCTTC | GCTTTCGCCATTTGGCTCTCCTATT | AA | ATCATCCAGTAAACCGCC |
| 12 | CCTCGTAAATCCTCATCA | AA | TCGAGTAAACACATCCGGATAATGA | TAGATTAACTCTCATCGCCAATTCC | AA | ATCATCCAGTAAACCGCC |
| 13 | CCTCGTAAATCCTCATCA | AA | AGTTGAAAGCTGCTGAATGTGGTCC | CTTCCGAAGACCTTCTCCAGTTCTT | AA | ATCATCCAGTAAACCGCC |
| 14 | CCTCGTAAATCCTCATCA | AA | AATCGTCGAGATGATCGGGTGACGG | ACCTCCTCTGCTTCCTTTTTGGCGG | AA | ATCATCCAGTAAACCGCC |
| 15 | CCTCGTAAATCCTCATCA | AA | CGCGATTGACGACATCTTATGTTGG | GCCGTCCGATAATCGGCGATGCATT | AA | ATCATCCAGTAAACCGCC |
| 16 | CCTCGTAAATCCTCATCA | AA | ACGTCAACATCGTCGGTTACGGGTT | GATCGGTCGGTTTCGTCGCAGGTTA | AA | ATCATCCAGTAAACCGCC |
| 17 | CCTCGTAAATCCTCATCA | AA | CGGCTCCATAAAATATGATATTCCC | ACGGCGGGCGGATGGAGTTAGATTA | AA | ATCATCCAGTAAACCGCC |
| 18 | CCTCGTAAATCCTCATCA | AA | AGACAATGAGATCTGTAACAGTTAC | TAATTAATCTTGATGACGAGTTCCC | AA | ATCATCCAGTAAACCGCC |
| 19 | CCTCGTAAATCCTCATCA | AA | AACTCGTCTTCACGTGACGGTTAAC | GTTGTTGTATCGGATAATCGCGCGA | AA | ATCATCCAGTAAACCGCC |

**Table S10.** Primer pairs for genes targeted in *Phalangium opilio* RNAi experiments and *Parasteatoda tepidariorum* colorimentric *in situ* hybridizations.

| **Species** | **Gene name** | **Primer name** | **Primer sequence** | **Amplicon length** |
| --- | --- | --- | --- | --- |
| *Phalangium opilio* | *clawless* | Popi_cll_for | GCAACAGCTGAACGAACTCA | 812 bp |
| *Phalangium opilio* |  | Popi_cll_rev | TATACGTGGGTGGTCGATCG |  |
| *Parasteatoda tepidariorum* | *clawlessA* | Ptep_cllA_for | TTTACTTTTGACATTTAATC | 1173 bp |
|  |  | Ptep_cllA_rev | CTAGCCCATGGTTGCAAAGT |  |
| *Parasteatoda tepidariorum* | *aristaless* | Ptep_al_T7_for | ggccgcggAGCTTCCTCGGTCTTGGTAA | 842 bp |
|  |  | Ptep_al_T7_rev | cccggggcTGGTTTTCCAGATCCTGGTG |  |
